# Fixation dynamics of beneficial alleles in prokaryotic polyploid chromosomes and plasmids

**DOI:** 10.1101/2022.01.07.475421

**Authors:** Mario Santer, Anne Kupczok, Tal Dagan, Hildegard Uecker

## Abstract

Theoretical population genetics has been mostly developed for sexually reproducing diploid and for monoploid (haploid) organisms, focusing on eukaryotes. The evolution of bacteria and archaea is often studied by models for the allele dynamics in monoploid populations. However, many prokaryotic organisms harbor multicopy replicons – chromosomes and plasmids – and theory for the allele dynamics in populations of polyploid prokaryotes remains lacking. Here we present a population genetics model for replicons with multiple copies in the cell. Using this model, we characterize the fixation process of a dominant beneficial mutation at two levels: the phenotype and the genotype. Our results show that, depending on the mode of replication and segregation, the fixation time of mutant phenotypes may precede the genotypic fixation time by many generations; we term this time interval the heterozygosity window. We furthermore derive concise analytical expressions for the occurrence and length of the heterozygosity window, showing that it emerges if the copy number is high and selection strong. Replicon ploidy thus allows for the maintenance of genetic variation following phenotypic adaptation and consequently for reversibility in adaptation to fluctuating environmental conditions.

## Introduction

Genetic variation is an important determinant of a population’s capacity to adapt to novel environmental conditions. In monoploid organisms, genetic variation exists only at the level of the population, whereas polyploid organisms may also be genetically heterogeneous at the intracellular level. In diploid eukaryotic organisms, observed heterozygosity – the carriage of different alleles by the two copies of a chromosome within a cell – is an important measure of genetic variation. In contrast, the existence and importance of intracellular genetic variation in prokaryotes has been so far much less appreciated; nonetheless polyploidy is common in prokaryotic species (Soppa, 2021). Polyploid chromosomes have been described across a wide range of taxa including cyanobacteria (Griese et al., 2011; Watanabe, 2020), gammaproteobacteria (Ionescu et al., 2017), as well as halophilic and methanogenic archaea (Breuert et al., 2006; Hildenbrand et al., 2011; Soppa, 2017). The number of chromosome copies in prokaryotes ranges from a few to several hundreds, and may also depend on the growth phase and the nutrient conditions (e.g., Maldonado et al., 1994; Hildenbrand et al., 2011; Watanabe, 2020). In bacterial species that are monoploid during slow growth, the number of chromosomes may temporarily increase during exponential growth (Nielsen et al., 2007; Sun et al., 2018). Indeed, early studies of bacterial genetics already observed heterozygosity in seemingly monoploid bacterial species such as *Escherichia coli* (Morse et al., 1956), *Bacillus subtilis* (Iyer, 1965), or *Streptococcus pneumoniae* (Guerrini and Fox, 1968). In addition to chromosomes, extrachromosomal genetic elements, such as bacterial plasmids, are often present in multiple copies in the cell. The plasmid copy number depends on the plasmid type and the environmental conditions, with some plasmid types reaching hundreds of plasmid copies in the cell (Friehs, 2004; Rodriguez-Beltran et al., 2021).

In sexually reproducing eukaryotes, heterozygosity is typically generated at the formation of zygotes. In prokaryotes, heterozygosity is generated through *de novo* mutations or recombination with DNA acquired through lateral transfer, e.g., via transformation, conjugation, or transduction. The subsequent maintenance of heterozygosity over time depends on the allele dynamics in the population. Two key determinants of allele dynamics in the population are the mode of replicon inheritance and the fitness effect of the mutant allele. Depending on the mode of replicon inheritance, daughter cells may be exact copies of the mother cell or differ in the distribution of alleles. In the latter case, segregation of the mutant allele may lead to the emergence of homozygous mutant cells. If the mutation is beneficial and survives stochastic loss while rare, the mutant allele will then ultimately fix in the population. Processes occurring at the intracellular level during cell division thus play an important role in the evolutionary dynamics of alleles in multicopy replicons and in their fixation processes and times.

The process of beneficial allele fixation plays a role in the rate of adaptation and the maintenance of variation. During the fixation process, both novel and wild-type alleles coexist in the population; once the beneficial allele has been fixed in the population, genetic variation at the allele locus is eliminated. Modelling allele fixation times has a long history in mathematical population genetics dating back to Kimura and Ohta (1969). Most existing models, however, focus on allele fixation in diploid sexually reproducing or in monoploid species. A recent modeling study on the evolutionary dynamics of alleles in multicopy plasmids suggests that the fixation times of alleles emerging in high-copy-number plasmids are longer than those of alleles emerging in low-copy number plasmids (Ilhan et al., 2019). Furthermore, Halleran et al. (2019) point out that random segregation of plasmid copies allows for allele fixation, while deterministic segregation hinders allele fixation. Both results clearly show that the allele dynamics on multicopy replicons are strongly influenced by the replicon properties. Yet, the effect of different replication and segregation modes on the fixation process, depending on the strength of selection, is still poorly understood.

Here we develop a mathematical framework to model the fixation process of beneficial alleles on multicopy replicons in asexual unicellular organisms. Our framework is germane to the evolutionary dynamics of alleles in polyploid prokaryotic chromosomes and in multicopy plasmids. We apply a classical population genetic model – the time-continuous Moran model with selection – and include various modes of replication and segregation of multicopy replicons. With this model, we investigate the dynamics of dominant beneficial alleles in the population. In the analysis, we follow the frequencies of heterozygous and homozygous mutant cells throughout the fixation process. Allele fixation in our model is defined at the levels of the cell phenotype and genotype, and the fixation times at both levels are compared. Fixation of the mutant phenotype implies phenotypic adaptation of the population. We describe that – if the two fixation times do not coincide – genetic variation still persists during the time between fixation of the phenotype and fixation of the genotype.

## The Model

We consider a population of bacteria (or other prokaryotes) with a constant number of *N* cells, each carrying *n* copies of a replicon (e.g., a multicopy (polyploid) chromosome or plasmid). We assume that there are two genetic variants of the replicon, carrying the *wild-type* and the *mutant* alleles respectively. Consequently, for *n* > 1, cells might be either heterozygous (i.e., carrying both alleles) or homozygous (see Figure 1).

**Figure 1:**
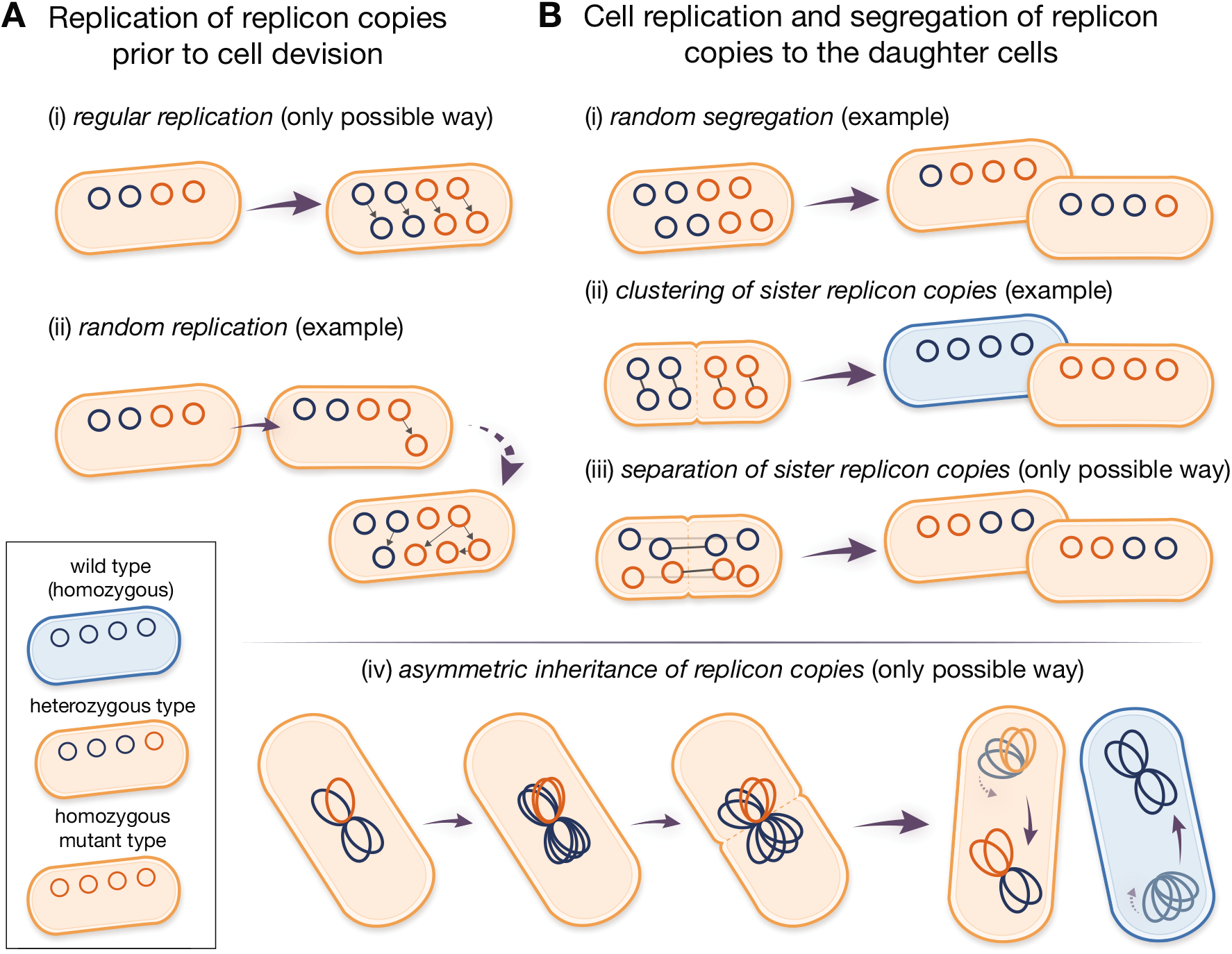
Modes of replication (A) and segregation (B) of the replicon copies modeled here. Blue and orange circles denote wild-type and mutant replicon copies respectively. Small gray arrows between replicon copies indicate sister replicons, i.e., one replicon copy is a direct duplicate of the other. (A) In case of regular replication, all replicon copies are duplicated exactly once before cell division. For random replication, replication of copies is a successive process. Random replication can lead to many different compositions of the replicon pool before cell division. (B) Given the mode of random segregation, different types of daughter cells can emerge. For *clustering of sister replicon copies,* all pairs of sister replicons (gray arrows) consisting of a template and its direct duplicate are inherited to the same daughter cell. The opposite holds for *separation of sister replicon copies,* where the composition of both daughter cells is identical due to partitioning of sister copies. In case of (iv) *asymmetric inheritance of replicon copies,* clusters of replicon copies are resolved at the deepest point in their genealogy ((iv) adapted from Sun et al. (2018)).

We assume that cells carrying at least one mutant replicon copy have a selective advantage *s* > 0. The mutation is thus dominant. Initially at *t* = 0, the mutant type is present in a single replicon copy in a small fraction *f* of cells. The initial population composition, as defined here, may arise, for example, due to a transformation event in the laboratory (or plasmid invasion). Later, we also consider adaptation starting from a balance between recurrent transformation and purifying negative selection against mutant cells that has established in a different environment, in which the mutation is deleterious. We do not consider the emergence of *de novo* mutations during the fixation process.

To describe the allele dynamics, we apply a classical population genetics model, the Moran model in continuous time (Moran, 1958), extended for multicopy replicons (Santer and Uecker, 2020). Mutant cells with *i* > 0 mutant replicon copies divide at rate λ_*i*_ ≡ 1 + *s*, while wild-type cells divide at rate λ_0_ ≡ 1. A dividing cell (parental cell) gives rise to two daughter cells, which replace the parental cell and one additional, randomly chosen, cell in the population. The formation of *n* new replicon copies occurs in the model prior to cell division. Here, we consider two modes of replication: regular replication and random replication (Figure 1A, see also Novick and Hoppensteadt, 1978). In the regular replication mode, each replicon copy is duplicated prior to cell division. This is assumed for the replication of many chromosome types (Skarstad et al., 1986; Nordström and Dasgupta, 2006). In the random replication mode, the following procedure is repeated *n* times: a single replicon copy is randomly selected for replication and the replicated copy is added to the replicon pool until the total number of replicon copies is 2*n* in the cell. This mode better reflects the replication mechanism of plasmids (Rownd, 1969; Bogan et al., 2001; Nordström, 2006). Possibly but not necessarily, it might also reflect the replication of some polyploid chromosomes as in some cyanobacteria, where only few chromosome copies are duplicated at once (Ohbayashi et al., 2019; Watanabe et al., 2012; Soppa, 2021). At cell division, the total replicon pool is divided equally between the daughter cells, i.e., each daughter cell receives *n* copies. In our baseline model, we assume that the segregation of mutant and wild-type replicons to the daughter cells is random (Figure 1B(i)). Mathematically speaking: n copies are drawn from the pool of 2*n* replicon copies of the parental cell without replacement and segregate to the first daughter cell; the remaining n copies are segregated to the second daughter cell. Chromosome segregation is random or at least partially so in a range of bacterial and euryarchaeotic species (Hu et al., 2007; Schneider et al., 2007; Tobiason and Seifert, 2010; Li, 2019). This mode moreover mimics the segregation of high-copy number plasmids (Ishii et al., 1978; Novick and Hoppensteadt, 1978; Cullum and Broda, 1979). Note that randomness in segregation in our model refers to the random segregation of replicon variants. Yet, segregation of high-copy number plasmids includes in addition randomness in the number of copies that each daughter cell inherits (Münch et al., 2015). Likewise, active partitioning systems in low-copy number plasmids may only guarantee that no plasmid-free cells are generated but do not necessarily imply equal plasmid copy numbers in both daughter cells following cell division. We simplify this in our model to keep the number of cell types manageable.

In addition to the baseline model, we consider three further modes of segregation: (ii) *clustering of sister replicon copies*, (iii) *separation of sister replicon copies*, and (iv) *asymmetric inheritance of replicon copies* (Figure 1B). Sister replicon copies are pairs where one copy is the direct replicate of the other. We only consider those in combination with regular replication, which is in some cases biologically motivated (e.g., for mode (iv)) and in others mathematical convenience (e.g., for mode (iii)).

In the segregation mode termed *clustering of sister replicon copies* (ii), sister replicons are inherited to the same daughter cell, while in the segregation mode termed *separation of sister replicon copies* (iii), the sister replicons segregate into different daughter cells. *Clustering of sister replicon copies* may happen in the presence of DNA binding regulatory elements (Wu et al., 1992), which has been recently shown to affect plasmid allele segregation under non-selective conditions (Garoña et al., 2021). It could also serve as a rough proxy of chromosome segregation when chromosome copies are spatially sorted in the cell as in *Synechococcus elongatus* (Jain et al., 2012). In this mode (ii), we only consider even copy numbers *n* to be able to fulfill the assumption of equal copy numbers in both daughter cells after cell division. The separation of sister replicon copies (iii) assumes that sister replicons are well separated post replication, as recently shown for haploid *Bacillus subtilis* chromosomes (Wang et al., 2017). The replicon separation may be achieved by active partition systems that push the replicons to the opposite cell poles such that they end up in different daughter cells at cell division, which is encoded in many low-copy-number plasmids (Nordstrñm and Gerdes, 2003; Million-Weaver and Camps, 2014; Brooks and Hwang, 2017). *Asymmetric inheritance of replicon copies* (iv) has been proposed by Sun et al. (2018) as a model for segregation of chromosomes in fast-growing bacteria, which harbor multiple chromosome copies due to multifork replication (Nielsen et al., 2007; Sun et al., 2018). Here the replicon copy number n is restricted to powers of 2. In this mode, all replicon copies remain attached to each other and form one large cluster. At cell division, only the oldest link between the replicon copies is resolved so that n copies are inherited to every daughter cell. Effectively, this means that one of the daughter cells of a heterozygous progeny cell receives all mutant copies. A mathematical description of the model is given in section A.1.

In our model, we track the fraction of cells carrying *i* mutant replicon copies over time *t*, which we denote by *x_i_*(*t*). A time unit corresponds to the mean generation time of wild-type cells. For most of the analysis, we study the deterministic dynamics, which are given by a system of n +1 ordinary differential equations (Eq. (A.11), Appendix A.2). We numerically integrate these equations using the Python package SciPy (Function `solve_ivp`). We determine the proportion of heterozygous cells *x*_het_ ≡ *x*_1_ + … + *x*_*n*-1_ and the proportion of homozygous mutant cells *x_n_* for all times *t*.

### Data availability

The authors state that all data necessary for confirming the conclusions presented in the article are represented fully within the article. Supplementary information and figures can be found in File S1. The simulation code and the scripts used for computer algebra are stored in File S2.

## Results

To describe the fixation dynamics, the population-wide frequency of the mutant replicon is reported at two levels: the phenotype level and the genotype level. Since we consider a dominant mutant allele, all cells that carry at least one mutant replicon copy have the same phenotype (i.e., fitness in our context). Fixation at the phenotype level is never strictly reached since new wild-type cells are constantly regenerated at divisions of heterozygous cells. Let us ignore this for a moment and denote the time by which (nearly) all cells contain at least one mutant replicon copy by *t*_phen_. From the time point of mutant phenotype fixation, *t*_phen_, selection is mostly restricted to the dynamics of wild-type homozygous cells that are newly generated at cell division of heterozygous cells and their few progenitors. The allele segregation process, followed by purging of wild-time homozygotes, continues until all cells have lost the wild-type replicon variant, i.e., the population is entirely composed of homozygous mutant cells. At time *t*_fix_, the mutation is fixed at the genotype level, and the wild-type variant has been lost from the population. In a deterministic model, true fixation never occurs, and we define *t*_phen_ as the time by which 99% of cells contain at least one mutant replicon copy and *t*_fix_ as the time by which 99% of cells are mutant homozygotes.

### Phenotypic and genotypic fixation times can differ for multicopy replicons, leading to a ‘heterozygosity window’

Notably, fixation of the mutant allele at the genotype level can occur a long time after its fixation at the phenotype level; here, we term the time interval between these two events the *heterozygosity window* (Figure 2). The length Δ*t* = *t*_fix_ – *t*_phen_ of the heterozygosity window is important since, during this phase, the population is fully adapted; yet, genetic variation is preserved. This may enable the population to quickly adapt if the selection pressure is reversed and the wild type becomes beneficial, although the potential to readily respond to this new change will ultimately also depend on the dominance or recessiveness of the wild-type allele under the reversed conditions. The total time during which genetic variation persists in the population, either within cells (heterozygosity) or across cells, is given by the genotypic fixation time.

**Figure 2:**
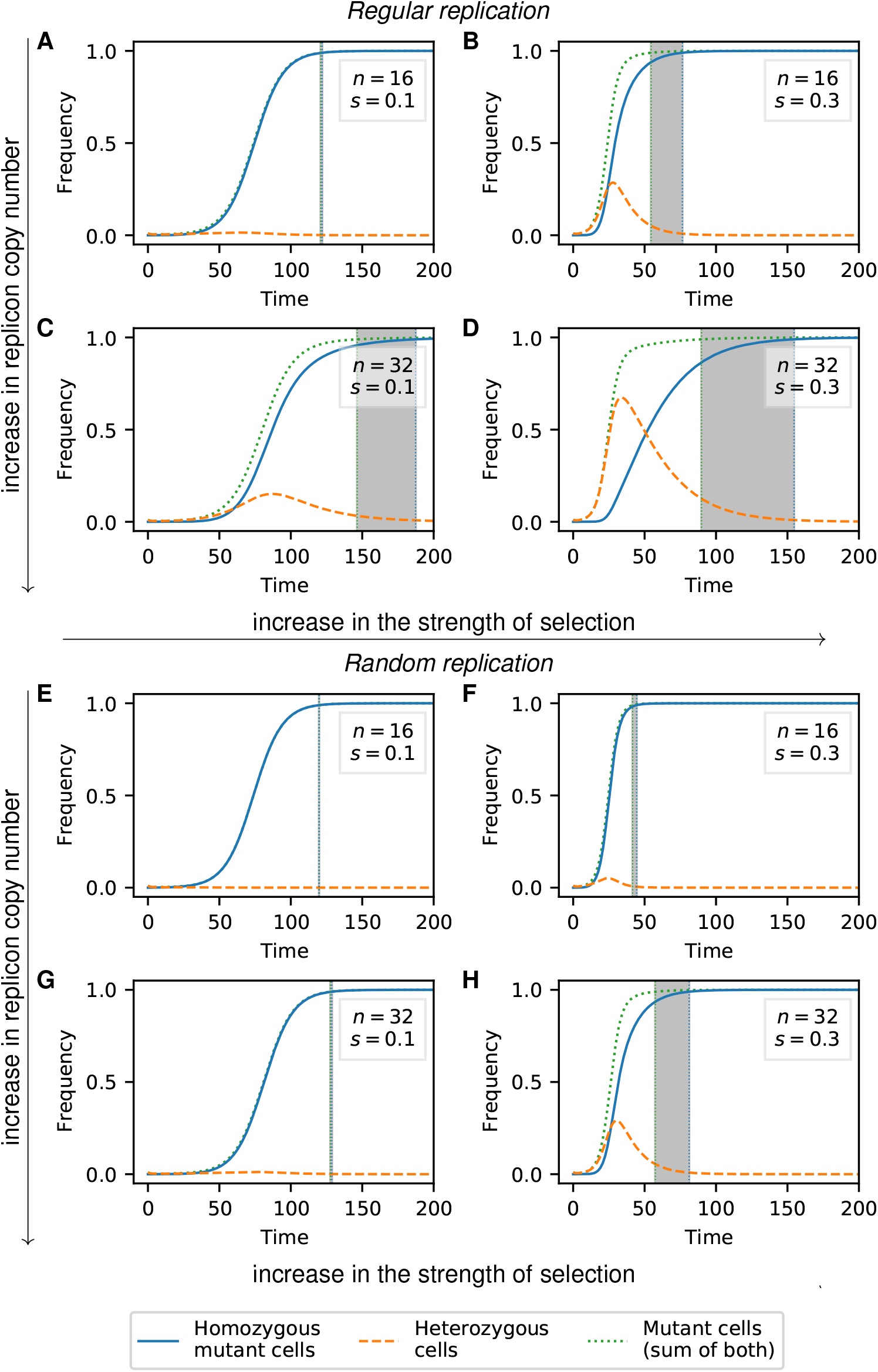
Frequency trajectories of different cell types for random segregation and regular replication (Panels A-D) or random replication (Panels E-H). The initial population at *t* = 0 comprises a small fraction *f* = 0.01 of heterozygous cells with one mutant replicon copy each. The gray area highlights the heterozygosity window, defined as the time between fixation of the mutation at the phenotype level (99% of cells carry at least one mutant copy) and fixation at the genotype level (99% of cells are mutant homozygotes). Results for cell frequencies were obtained from the deterministic model in Eq. (A.11).

We find that a heterozygosity window appears if the replicon copy number n and the strength of selection s are sufficiently large (Figures 2, 3). If n and s are large, heterozygous cells rise considerably in frequency before homozygous mutant cells become frequent and take over the population. We can use this insight to derive a condition for the existence of a heterozygosity window. We find that for regular replication and small initial mutant frequencies *f*, heterozygous cells initially increase in frequency if

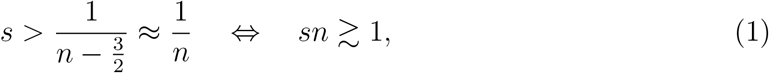

where the last approximation holds for 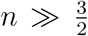 (see A.3 for a mathematical derivation). This condition predicts well the boundary in the s-n plane between areas with and without a heterozygosity window (Figures 3B, S2A).

**Figure 3:**
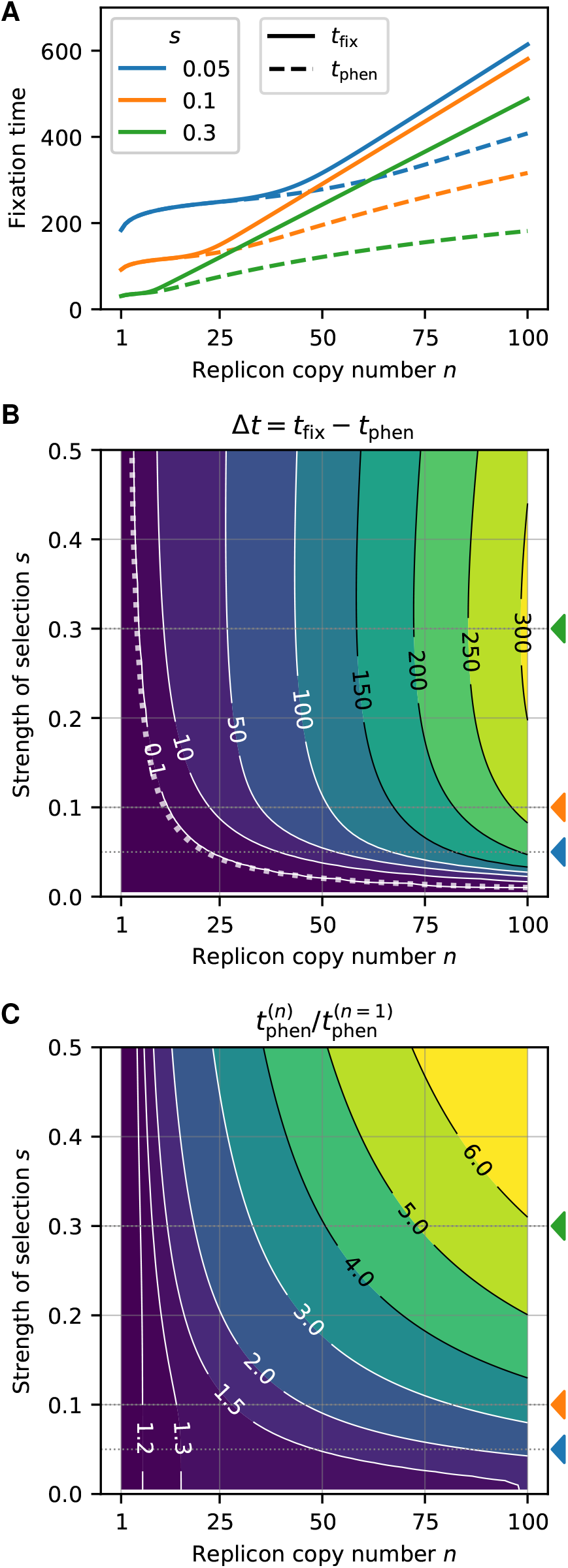
Effect of the replicon copy number *n* and the strength of selection *s* on the fixation times and the heterozygosity window for a replicon subject to regular replication and random segregation. The initial frequency of mutant cells with one mutant replicon copy at *t* = 0 is *f* = 0.01. (A) Fixation times as a function of the replicon copy number for several selection coefficients *s* = 0.05 (blue), 0.1 (orange), 0.3 (green) (indicated by colored triangles in Panels B and C). Lines are for guidance of the eye; the replicon copy number n is discrete. (B) Contour plot of the heterozygosity window for various replicon copy numbers n and selection coefficients s. The dotted line shows the threshold of s (as a function of *n*) at which the heterozygosity window starts to occur (criterion (1)). (C) Time of fixation at the phenotype level *t*_phen_ relative to *n* = 1. All graphs show results of deterministic numerical simulations (Eq (A.11)). Fixation times *t*_phen_ and *t*_fix_ were determined as the I time point when mutant and homozygous mutant cells reach a threshold of 0.99 respectively.

If multicopy replicons undergo random rather then regular replication, the threshold of s and *n* for a heterozygosity window to appear is higher. Analogous to Eq. (1), we find the condition

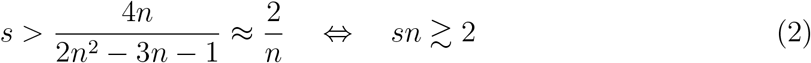

(Figures 2 and S2B), i.e., the strength of selection needs to be twice as strong or the copy number twice as large for a heterozygosity window to appear. This is consistent with the finding that the decay of heterozygotes through replicon segregation is faster under random replication than under regular replication, where the heterozygote loss rates are 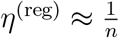 and 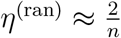 for regular and random segregation, respectively (Eqs. (A.16) and (A.19)).

With our model, the choice of the fixation threshold can influence the length of the heterozygosity window Δ*t* (*x*_thr_ = 99% in all results in the main text). If the initial frequency of mutant cells is very small, *f* ≪ 1%, the heterozygosity window length Δ*t* is smaller with *x*_thr_ = 99% than with a larger threshold (see Figure S1 for an example). In the limit *x*_thr_ → 1, however, it converges to a size that is independent of the specific choice of f (see Figure S1 and supplementary information section S1 for a mathematical proof). This shows that the appearance of the heterozygosity window is a robust phenomenon and not an artifact of our specific choices of *f* and *x*_thr_.

### The heterozygosity window is large if the copy number is high and the selection strong

The fixation times at the phenotype and genotype levels *t*_phen_ and *t*_fix_ both increase with the replicon copy number (Figure 3A). This is not a consequence of our choice of the initial condition, for which the initial frequency of mutant replicon copies *f*_rep_ = *f/n* is smaller for higher n: keeping *f*_rep_ rather than f constant, the fixation times are independent of n for small n and s where no heterozygosity window occurs but still increase with n otherwise (Figure S4).

The heterozygosity window length Δ*t* increases with the replicon copy number n and – over large parts of the parameter range – with the strength of selection s (Figures 2, 3B, S5B). For very high strength of selection s, the heterozygosity window length Δt is again smaller due to a decrease in the overall fixation times. If scaled with the fixation time *t*_fix_, the size of the heterozygosity window Δ*t/t*_fix_ monotonically increases with s and eventually converges (Figure S3).

A mathematical analysis of the fixation process (provided in File S1) shows that the het-erozygosity window length is approximately given by

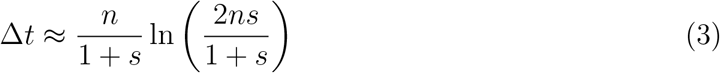

for regular replication and by

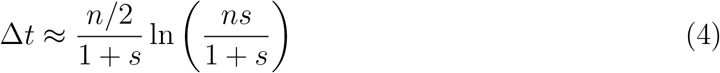

for random replication. Hence, the implementation of random replication in the model leads to a reduced window size in a similar manner as a reduction in the replicon copy number by a factor of 2.

### The heterozygosity window also exists in small finite populations but is smaller

The deterministic analysis in the previous section ignores stochastic fluctuations in the genotype frequencies, reflecting the dynamics in an infinite or very large population. To account for finite population sizes, we complemented our analysis with stochastic simulations. Unlike in the deterministic model, the mutant allele can go extinct while rare, and we consider fixation times conditioned on fixation of the mutant allele. To render the results comparable to those of the deterministic model, we again define that phenotypic fixation is reached when 99% of cells are mutant. Similarly, fixation at the genotype level is reached when 99% of cells are homozygous mutant.

We find that a heterozygosity window also occurs in finite populations (Figure 4). For small populations, the heterozygosity window is smaller than predicted by the deterministic model, especially if the replicon copy number *n* is high (Figures 4C and D, S6).

**Figure 4:**
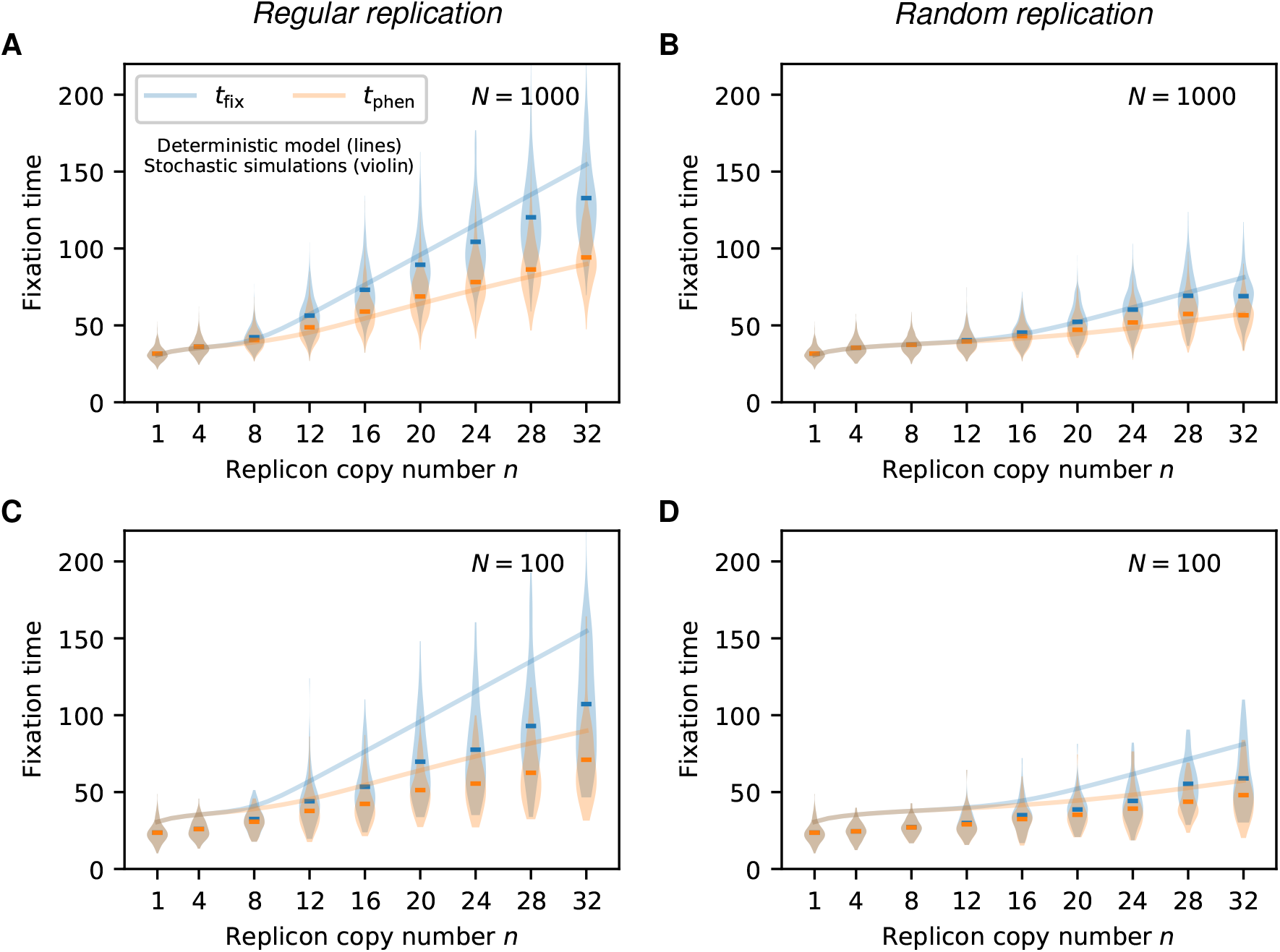
Fixation times *t*_phen_ and *t*_fix_ from stochastic simulations for selected replicon copy numbers (violin plots, mean shown as horizontal line) in comparison to results from the deterministic model (lines), *s* = 0.3. To compare between stochastic and deterministic calculations, we define fixation at the phenotype level *t*_phen_ (resp. at the allele level *t*_fix_) by the state where the frequency of mutants (resp. of mutant homozygotes) hits a threshold of 0.99. For all 1000 stochastic trajectories, we determine the fixation time *t*_phen_ (*t*_fix_) by the mean of all time points at which the fixation threshold is reached.

For monoploid populations (*n* = 1), the expected fixation time of a mutant allele decreases with the population size (Kimura and Ohta, 1969). Similarly, we find that fixation of ho-mozygous mutant cells *t*_fix_ is faster in finite populations than predicted by the deterministic model; furthermore, the time to fixation decreases with the population size (Figure 4). The phenotypic fixation time *t*_phen_, however, reaches a maximum for an intermediate population size (cf. the fixation times for *N* = 1000 with those for *N* = 100 and those predicted by the deterministic model reflecting an infinite population in Figures 4A and S6).

### A heterozygosity window also exists if the cell-type frequencies are in trans-formation-selection balance prior to adaptation

So far, we assumed a given initial frequency *f* of mutant cells where each of those cells carries one mutant copy. This corresponds, for example, to the cell-type composition after incorporating a mutant allele into the plasmid via transformation (e.g., Garoña et al., 2021). In natural settings, however, mutations are often present at low levels for a long time in a balance between negative selection and recurrent appearance before they become beneficial due to a shift in the environmental conditions. In that case, cells with more than one mutant replicon copy may arise before the fixation process ensues.

In the following, we therefore model two phases – the first one, in which the mutant allele is subject to negative selection, modeled by a reduced cell division rate 1 – *σ* of mutant cells, and a second one, when it has turned beneficial and rises to fixation. For the first phase, we assume that the mutant allele appears in single replicon copies at a transformation rate *τ* per cell per time unit and determine the mutant cell frequencies in the equilibrium between the input of the mutant allele via transformation and loss due to negative purifying selection. At time point *t* = 0, the mutant allele becomes beneficial, i.e., mutant cells divide at rate 1 + *s* as in the above sections. A detailed description of the model is given in Appendix A.4.

Overall, we find that the general pattern of the heterozygosity window occurrence remains unchanged, irrespective of *τ* and *σ*: a window opens up for sufficiently large *n* (Figure 5).

**Figure 5:**
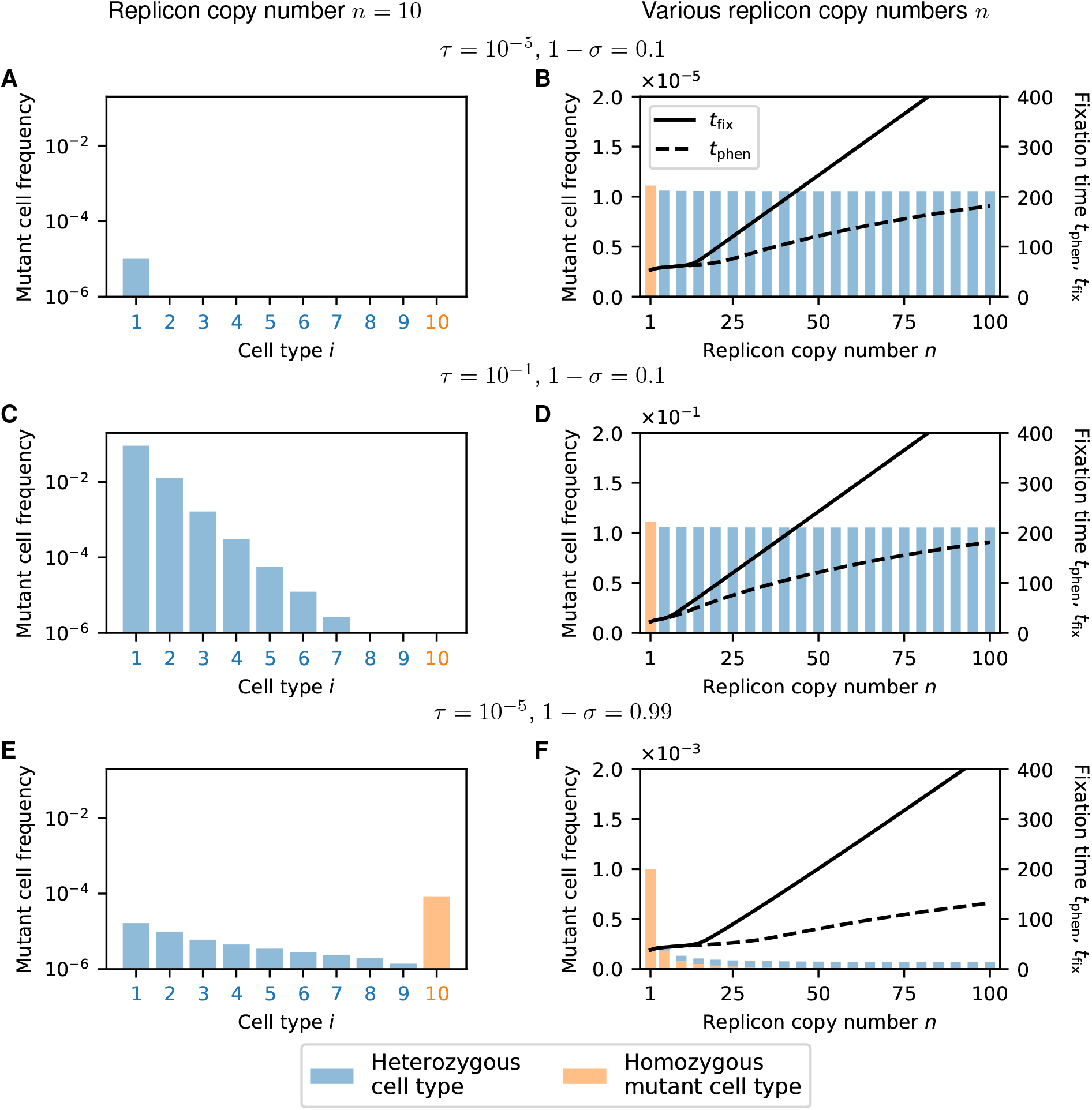
Mutant cell frequencies at transformation-selection balance for replicon copy number *n* =10 (A,C,E) and for various replicon copy numbers n (B,D,F). (A,C,E) Example with *n* = 10 replicon copies per cell. Deterministic frequencies of the different mutant cell types (number of mutant replicon copies per cell, *x*-axis) in the population. The remaining fraction of cells are wild-type homozygotes with *i* = 0. Single mutant replicon copies enter cells at a rate *τ*, and decrease the cell division rate by *σ*. Cell-type frequencies are shown on a logarithmic scale, allowing a direct comparison between various selection coefficients *σ* and transformation rates *τ*. (B,D,F) Mutant cell frequencies for various replicon copy numbers *n* (*x*-axis). Bar plots show mutant frequencies on a linear scale, which allows a comparison of the composition of heterozygous (blue) and homozygous mutant cells (orange). Note that the scale differs in all three panels. Line plots show times until fixation at the phenotype level tphen (dashed) and at the genotype level *t*_fix_ (solid) when the process is started at transformation-selection balance. Note that in Panel B, the heterozygosity window would open for smaller values of *n* if the thresholds for *t*_phen_ and *t_fix_* were chosen closer to 1. During the fixation process, mutant cells divide at rate 1 + *s* with *s* = 0.3 (wild-type cells divide at rate 1). Fixation times were obtained from deterministic simulations Eq. A.11) as in Figure 3.

Strongly deleterious mutations (cell division rate 1 – *σ* close to 0) mostly occur in heterozygous cells with few mutated replicon copies in transformation-selection balance (Figures 5A-D). Furthermore, the frequency of mutant cells is nearly independent of n (Figures 5B and D). For very low transformation rates, most of the mutant cells contain a single mutant replicon copy, which resembles the scenario that we considered in the above sections (Figure 5A). For high transformation rates, cells with more than one mutant replicon copy exist, which reduces fixation times for low-copy replicons but not for high-copy replicons (compare Figures 5B and D).

If the strength of selection is weak (1 – *σ* close to 1), mutant cells can persist longer in the population on average. Therefore homozygous mutant cells can be generated and exist at transformation-selection balance for low-copy numbers (Figures 5E and F). For high replicon copy numbers *n*, however, too many generations would be needed for homozygous cells to emerge; thus, almost all mutant cells are heterozygous at transformation-selection balance even if selection is weak. Unlike for strong selection, the overall frequencies of mutant cells strongly decrease with *n*. Nonetheless, for high *n*, fixation times are smaller compared to the case of strong selection (Figure 5F and B).

### The mode of replicon segregation strongly influences the occurrence and length of a heterozygosity window

In the previous sections, we considered random segregation of replicon copies at cell division. Here, we examine the effect of alternative segregation modes on the fixation dynamics (Figure 1). All three alternative modes in our model reflect a more deterministic form of replicon inheritance compared to the baseline model of random segregation.

Notably, the *clustering of sister replicon copies* segregation mode reduces the unit of inheritance – that is, the number of segregating DNA molecules – by a factor of two compared to random segregation (1). The fixation dynamics under *clustering of sister replicon copies* with copy number 2*n*, therefore, resembles the resulting dynamics under random segregation with replicon copy number *n*. Both the fixation time of mutant cells and of homozygous mutant cells are reduced and the heterozygosity window is smaller if sister replicons segregate into the same cell than if they segregate independently from each other (Figures 6A and S7A, cf. 2D and 3A for the baseline model with random segregation). In line with our other results on random and regular replication (Eq. 3 and 4), fixation times *t*_p_h_en_ and *t*_fix_ and the size of the heterozygosity window for *clustering of sister replicon copies* are very similar to those obtained for random replication (cf. Figures S5A and S7A).

**Figure 6:**
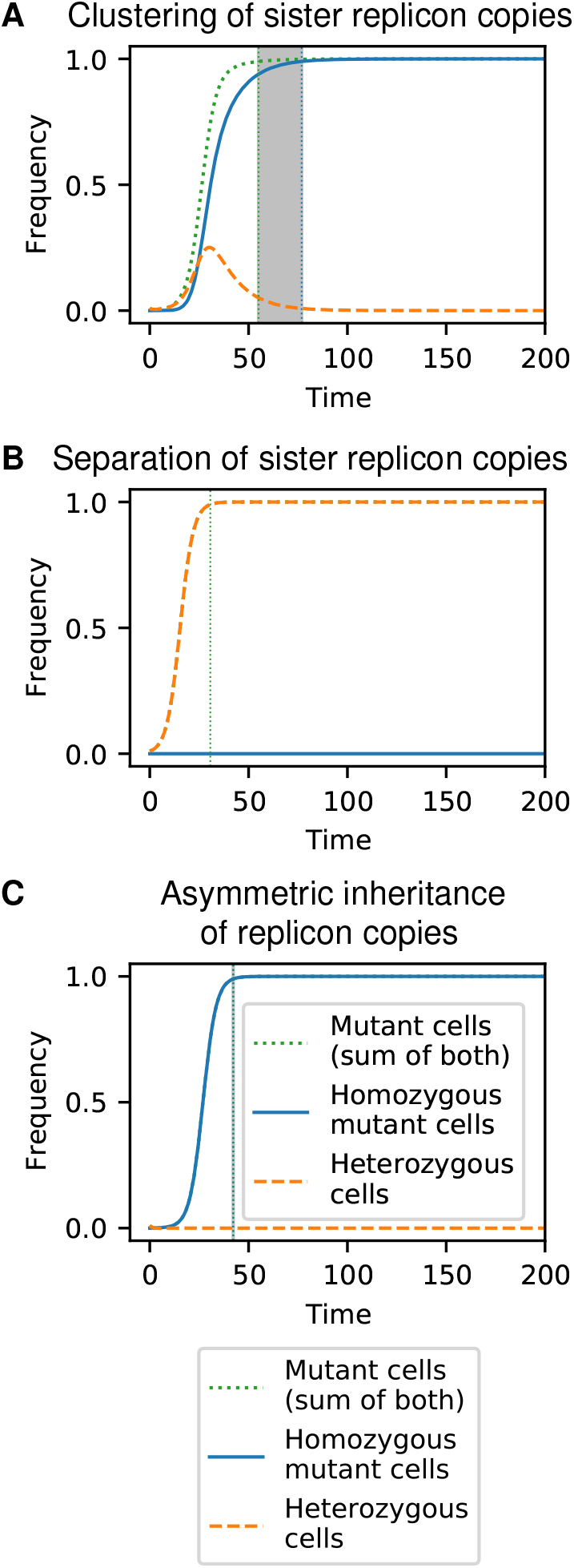
Frequency trajectories of different cell types for alternative models of replicon segregation in combination with regular replication: For (A) *clustering of sister replicon copies,* all pairs of sister replicon copies are inherited to the same daughter cell, whereas for (B) *separation of sister replicon copies,* sister replicon copies segregate into different daughter cells. (C) For the mode of *asymmetric inheritance of replicon copies,* one of the daughter cells of a heterozygous cell reveives all mutant replicon copies. Parameters: Replicon copy number *n* = 32, strength of selection *s* = 0.3, initial frequency of mutant cells with one mutant replicon copy *f* = 0.01 (cf. Fig. 2D for the baseline model of random segregation). Calculations and visualization were performed as in Figure 2.

Under the mode of *separation of sister replicon copies*, cells with *i* mutant replicon copies always produce two daughter cells of the same type *i* since every sister couple is equally divided between the two daughter cells. Hence, the mutation will never reach fixation at the genotype level (Figure 6B), and in the absence of gene conversion and without deviations from the model, heterozygosity is maintained forever. The fixation dynamics at the phenotypic level are effectively reduced to the case *n* = 1 as wild-type cells divide into wild-type cells, and mutant cells divide into mutant cells of the same type *i* (Figure S7B). Consequently, the fixation time of mutant cells is independent of the replicon copy number in the *separation of sister replicon copies* mode.

Last, we consider the *asymmetric inheritance of replicon copies* mode, which reflects a more extreme scenario of sister replicon clustering. Following the mode of asymmetric inheritance, heterozygous cells always divide into one heterozygote and one homozygous cell. Consequently, there is no increase in heterozygous cells, and no heterozygosity window appears (Figure 6C). A comparison of the fixation dynamics for different replicon copy numbers shows that the fixation time increases slightly with the copy number n (Figure S7C). The reason for this increase is the smaller initial mutant replicon frequency on the allele level *f*_rep_ = *f/n* for higher *n*. For a constant initial mutant frequency on the allele level *f*_rep_ = *f/n*, the fixation time is independent of the copy number n in this inheritance mode (cf. Figure S8).

## Discussion

To understand the consequences of polyploidy for allele dynamics in prokaryotes, we considered the fixation process of a dominant beneficial mutation on a multicopy replicon.

### Maintenance of heterozygosity on multicopy replicons

Our initial model in which replication is regular and segregation random shows that fixation times are longer on multicopy replicons than on single-copy replicons and increase with the copy number. This is generally in line with experimental and theoretical results by Ilhan et al. (2019), who simulated the distribution of replicon copy variants in the daughter cells and the cell composition in the next generation by binomial sampling. For large copy numbers and strong selection (see Eq. 1), we moreover find a delay between fixation at the phenotype level and fixation at the genotype level, which we term ‘heterozygosity window’. Within the heterozygosity window, the population is phenotypically fully adapted, while genetic variation is maintained. For example, *de novo* evolution of antibiotic resistance would be reversible during the heterozygosity window if antibiotics are removed, and the resistance mutation has a negative fitness effect in the absence of antibiotics. Importantly, such a reversible adapation would leave no trace in the genome. However, how easily the population can adapt to such a future change also depends on the dominance relationship between the two alleles in the new environment (e.g., antibiotic-free environment), and future models are needed to assess this. In our model, no heterozygosity window emerges if selection is weak or the copy number is low. These results hold, irrespective of whether the adaptive process starts from a constant low fraction of mutant cells with one mutant copy each, a constant low fraction of mutant replicon copies (with no more than one copy in each cell), or from transformation-selection balance (Fig. 3A and Fig. 5).

The existence of a heterozygosity window can be understood in the following way: if selection is strong, heterozygous cells quickly rise in frequency. At the same time, if the copy number is large, homozygous cells emerge only slowly, and the mutant cells become fixed before mutant homozygotes dominate the population. From this point on, the process is selectively neutral except for homozygous wild types that are generated during cell division. The emergence of such wild-type cells is rare for high replicon copy numbers such that the fixation of homozygous mutant cells is slow. If selection is weak or the copy number low, homozygous mutant cells are generated early in the adaptive process. They quickly rise in frequency since all daughter cells of homozygous cells are themselves homozygous. Heterozygous cells, in contrast, rise only little or not at all since they segregate too many offspring into the homozygous classes. In that scenario, phenotypic fixation coincides with genotypic fixation.

Most of our analysis relies on a deterministic model for the genotype frequencies in the population. Stochastic simulations show that the heterozygosity window is smaller than predicted by the deterministic model if the population size is small, but qualitatively, the results also hold in small finite populations. Additionally, we find that the phenotypic fixation time has a maximum for intermediate population sizes.

In the present study, we focused on dominant mutant alleles, where a heterozygosity window is expected to be most prominent. For a recessive mutant allele, heterozygous cells have no selective advantage over wild-type cells, and therefore do not rise in frequency by natural selection. Once homozygous mutant cells finally emerge, they rapidly rise to fixation. The dynamics of mutant alleles of intermediate dominance (i.e., the cell fitness increases with the frequency of the mutant allele) are positioned between these two extremes, and the effects leading to a heterozygosity window would be less pronounced than for a dominant mutation. This is likely similar for alleles with a gene dosage effect (i.e., where the phenotype depends on the number of mutant replicon copies).

Experimentally, an initial rise and subsequent decline of heterozygotes and a heterozygosity window have been observed in invasion experiments of a beneficial allele on a multicopy plasmid (see Fig. 3 in Rodriguez-Beltran et al., 2018). Complementary computer simulations show that heterozygosity can be maintained for many generations if the selection pressures for the two alleles rapidly alternate. In these simulations, Rodriguez-Beltran et al. (2018) made the simplifying assumption that all heterozygous cells contain the two plasmid variants in equal proportions, i.e., there are only three cell types – the two homozygous types and heterozygous cells. Assuming that plasmid copies segregate to the daughter cells with probability 1/2, the probability that a heterozygous cell forms two homozygous cells at cell division is 2^1–*n*^ (using our model formulation, where plasmid copies are replicated prior to cell division). Using this assumption, 2^1–*n*^ is the heterozygote loss rate in the absence of selection (see A.3, Eq. (A.16)). Our analysis, which explicitly considers heterozygous cells with different compositions of the replicon pool, shows that the heterozygote loss rate is more accurately described by 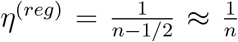 for regular replication (see Eq. (A.16)). Comparing these two results shows that the approximation in Rodriguez-Beltran et al. (2018) underestimates the loss rate, especially for high replicon copy numbers *n*. Similar to our model, Novick and Hoppensteadt (1978) calculated the decrease in the proportion of heterozygous cells per generation in a geometrically growing population as 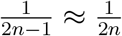 for plasmid copies undergoing regular replication. The difference of a factor 1/2 between the loss rate in our model and the loss rate in Novick and Hoppensteadt (1978) is due to the different population dynamics in the two models (constant population size vs geometric growth).

It is interesting to compare fixation on a multicopy replicon in an asexually reproducing population to the fixation of a beneficial allele in a diploid sexually reproducing population: in the latter, there is also a heterozygosity window for dominant (but not for recessive) alleles (Fig. S9). The underlying dynamics are, however, very different since the generation of homozygous individuals requires mating of heterozygous individuals, and mating of the two homozygous types re-generates heterozygous individuals.

### The effects of the segregation and replication modes

The modes of replication and segregation depend on the respective replicon type and on the species. In our study, we consider several fundamental modes for both processes. In the future, the model can be tailored to accurately describe the details of specific systems. For most of our analysis, we assumed that each replicon copy is replicated exactly once prior to cell division (regular replication). We considered four modes of replicon copy segregation. In all modes considered, both daughter cells inherit the same number of replicon copies, but the segregation mode affects the allele distribution in the daughter cells and hence the maintenance or loss of intracellular genetic variation. Under the mode *separation of sister replicon copies*, the heterozygosity window is infinitely long, i.e., heterozygosity is maintained forever. The complete opposite dynamics occurs under the mode of *asymmetric inheritance of replicon copies* where one of the daughter cells inherits the maximum possible number of mutant copies. In that case, heterozygous cells do not increase in frequency since cell division of heterozygotes leads to one heterozygous cell and one homozygous cell. Thus, in monoploid bacteria that become effectively polyploid during fast growth due to multifork replication, heterozygosity will rapidly decrease, and no heterozygosity window arises. The results obtained applying the modes of *random segregation* and of *clustering of sister replicon copies* lie between perfect separation of copies and asymmetric inheritance. In these modes, heterozygous subpopulations can rise transiently, given that the replicon copy number and the strength of selection are sufficiently high, and a heterozygosity window opens up. Since the replication of plasmids and likely also of some types of chromosomes is better described by random than by regular replication of copies, we also modeled the replication mode *random replication* (in combination with random segregation of replicon copies). In that case, the heterozygosity window is also present but smaller than under regular replication. Notably, our results show that it is approximately as large as for a regularly replicating replicon with a copy number *n*/2. E.g., for a plasmid with copy number *n* ≈ 20 that is undergoing random replication and segregation, there is only a delay of a few (wild-type) generations between phenotypic and genotypic fixation even if selection for the mutant allele is strong (Fig. S5). For a replicon undergoing regular replication, there would be a delay of around 40 generations with *s* = 0.3 (Fig. 3).

Our modeling framework could be applied to support experimental studies in polyploid species. The mode of chromosome segregation differs between prokaryotic species and is not always well understood. Following the fate of heterozygous cells has been used as one approach to gain insights into the segregation patterns (e.g., Pulakat et al., 1998; Suh et al., 2000; Tobiason and Seifert, 2010; Li, 2019). With our modeling framework, we can make quantitative predictions on the maintenance of heterozygosity and the time to loss or fixation of a marker. This makes it possible to test on a quantitative basis which segregation patterns are compatible with experimental observations and thus to better understand which conclusions can and cannot be drawn. A second application of our model concerns genetic engineering. Genetic engineering is known to be difficult in highly polyploid species such as *Synechocystis PCC 6803*, which carries approximately 60 chromosome copies per cell (Griese et al., 2011). To incorporate an allele into all chromosome copies, positive selection for this allele needs to be applied for a large number of generations. If selection is released too early, reversion to the wild type may occur. Our model allows us to estimate the required number of generations in advance. There is, however, a caveat with applying our current model to experimental studies: we here assume a constant population size, while in most experiments, the population size drastically changes in the alternation between exponential growth and population bottlenecks. This very likely affects the size of the heterozygosity window. Yet, our model can be readily adjusted to account for such population dynamics, e.g., by replacing the Moran model by a birth-death model and including bottlenecks at regular intervals.

In this study, we focused on the dynamics of multicopy replicon copies in prokaryotes. Similar dynamics and questions arise for polyploid eukaryotic cells. Mitotic cell division leads to a division of sister replicon copies. Some replicon types in eukaryotes, however, do not undergo mitosis, for example, chromosomes in the somatic macronucleus of ciliates (Morgens and Cavalcanti, 2015), nuclear extra-chromosomal DNA (ecDNA) in tumor cells (Bailey et al., 2020), mitochondria and other organelles (Lightowlers et al., 1997; Stewart and Chinnery, 2015; Ramsey and Mandel, 2019). Specifically, heteroplasmy is known to occur in mitochondria, and the spread of mutations in mitochondrial DNA through replication of mutated copies and segregation at cell division within an organism or across generations is highly relevant in the context of disease development (Lawless et al., 2020). In all the above cases – macronucleus of ciliates, ecDNA, mitochondria and other organelles -, the segregation of replicon copies is random or at least partially so, and related modeling approaches have been applied to either study variation in the number of replicon copies or of genetic variants (e.g., Kimura, 1957; Morgens and Cavalcanti, 2015; Pichugin et al., 2019; Lawless et al., 2020). Our results could thus also be of interest for multicopy replicons in eukaryotes.

#### Conclusion

Heterozygosity is commonly considered in diploid sexually reproducing organisms. Prokaryotic cells can be heterozygous as well if they harbour a multicopy replicon, i.e., a polyploid chromosome or a multicopy plasmid. The present work demonstrates that heterozygosity of multicopy replicons, hence genetic variation, can be maintained for extended periods of time – the heterozygosity window – during the fixation process of a dominant beneficial mutation.

## Acknowledgements

The authors thank Florence Bansept and Arne Traulsen for helpful discussions. We thank Giddy Landan for critical comments on the manuscript. M.S. is a member of the International Max Planck Research School for Evolutionary Biology and gratefully acknowledges the benefits provided by the program.

## A Appendix

### A.1 Mathematical formulation of the model and stochastic com puter simulations

We describe the dynamics of the system by a state vector **N**(*t*) = (*N*_0_(*t*), *N*_1_(*t*),…, *N_n_*(*t*)), where *N_i_*(*t*) denotes the number of cells with *i* mutant replicon copies (‘cells of type *i*’). For all times *t*, the total number of cells 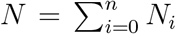 is constant, whereas the relative abundances of the different types may change. The number of cells of any type *i* can be altered either by cell division or by cell death. Cell death occurs by removal of a randomly chosen cell from the population right after cell division so that the total population size remains constant.

The rate at which a cell of type *i* divides into daughter cells of type (*j*_1_,*j*_2_) (ordered pair) is given by 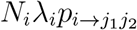, where *p*_*i*→*j*_1_*j*_2__ denotes the probability that cell division leads to daughter cells of type *j*_1_ (first daughter cell) and *j*_2_ (second daughter cell). The probability distribution *p*_*i*→*j*_1_*j*_2__ depends on the mode of replication and the mode of segregation and will be derived below for the various replication and segregation modes shown in Figure 1. The probability that a cell of type *l* is replaced following division of an *i*-type cell is given by *ν_l_* = *N/N* if *l* = *i* and *ν_l_* = (*N*_l_ – 1)/*N* if *l* = *i*. Thus, cell division events that increment the number of cells of type *j*_1_ and *j*_2_ (new daughter cells) and decrement the number of cells of type *i* (dividing cell) and of type *l* (replaced cell) occur at rate 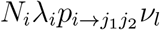. It should be noted that some or all of the cell types *j*_1_, *j*_2_,*i,l* may be identical. Cell division events that do not change the state vector **N** can be omitted in the simulations and when deriving the deterministic dynamics.

We now derive the probability distributions *p*_*i*→*j*_1_*j*_2__ for the different modes of replication and segregation. For *regular replication*, each replicon copy is duplicated, resulting in exactly *k* = 2_*i*_ mutated copies just before cell division of a type-i cell. For *random replication, k* is a random number. The successive replication of copies before cell division corresponds to a Pólya urn model (Eggenberger and Pólya, 1923; Mahmoud, 2008), and the probability to have k mutated copies before cell division is given by

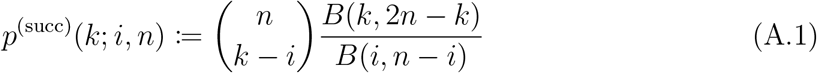

for *i* + *n* ≤ *k* ≤ *i* and zero otherwise, in case of heterozygous types 0 < *i* < *n*. *B* denotes the Beta-function, where, for positive integers,

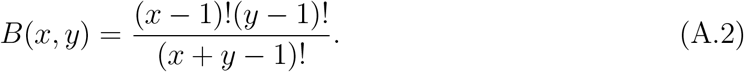

For homozygous types *i* = 0, *i* = *n*, we have *p*^(succ)^(*k; i,n*):= *δ*_*k*,2*i*_, where *δ*_*k*,2*i*_ denotes Kronecker’s delta.

Mutant and wild-type replicon copies are distributed to the daughter cells according to the chosen mode of segregation. In all of them, each daughter cell receives exactly *n* replicon copies in total. In the case of *random segregation*, the probability that a cell containing *k* mutant copies just before division produces two daughter cells of types (*j*_1_,*j*_2_) is given by

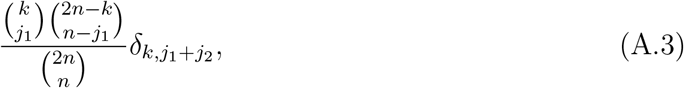

where *δ*_*k*;*j*_1_+*j*_2__ denotes Kronecker’s delta. If we combine this term with *k* = 2*i* for *regular replication* (reg) or with the probability distribution for *random replication* (ran), we obtain

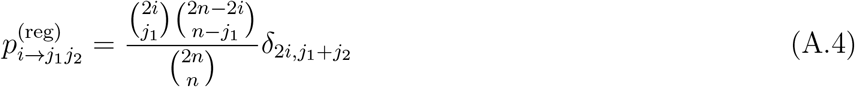

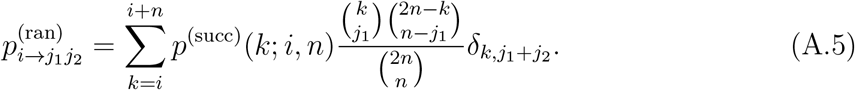

For the segregation modes, where sister replicons are clustered (ii) or separated (iii) to different daughter cells we only consider *regular replication* since *random replication* does not allow defining unique pairs of sister replicons. Similarly, we do not consider *random replication* for the segregation mode (iv) *asymmetric inheritance of replicon copies* since replicating one copy two times would violate our assumption of equal copy numbers in both daughter cells: For *asymmetric inheritance of replicon copies*, the two replicon copies that form the oldest link in genealogy are segregated to distinct daughter cells together with all their younger sisters (see below). Therefore, it is needed that each copy is duplicated (regular replication) once so that both daughter cells receive n replicon copies.

In the case of *clustered segregation of sister replicons* (clu) combined with *regular replication*, we need to consider the random distribution of *i* pairs of mutant replicon copies instead of 2*i* individual mutant replicon copies (cf. Equation A.4). The probability that an *i*-type cell with *i* pairs of mutant copies before cell division produces two daughter cells with (*j*_1_/2,*j*_2_/2) mutant couples respectively can be derived by replacing 2*i* → *i*, 2*n* → *n*, and 2*j_m_* → *j_m_*, *m* =1, 2 in Equation (A.4). We obtain

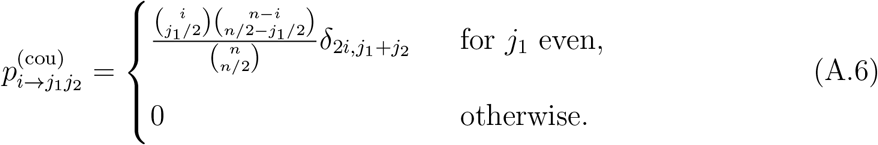

For this mode, we need to restrict the replicon copy number to even numbers *n*.

For the mode of *separation of sister replicon copies* (sep), reproduction of *i*-type cells produces only daughter cells of type *j*_1_ = *j*_2_ = *i*. Thus, we have

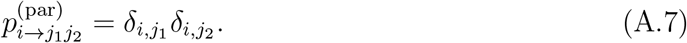

For the mode of *asymmetric inheritance of replicon copies* (asy), a heterozygous cell of type *i*, where 0 < *i* < *n*/2, divides into one daughter cell with twice the number of mutated copies than the parental cell and one daughter cell with only wild-type copies, i.e., *j* = 2*i* and *j*_2_ = 0. (From this, it follows that there are no heterozygous cells with *i* > *n*/2.) Homozygous cells divide into two homozygous daughter cells of the same type. Therefore, the probability that an *i*-type cell produces daughter cells (*j*_1_, *j*_2_) is given by

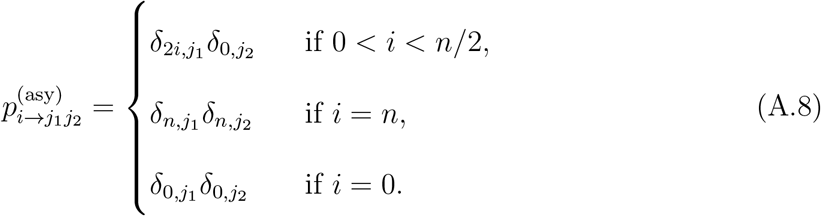

We perform stochastic computer simulations using the Gillespie algorithm (Gillespie, 1976), which implements the models exactly. The simulation code is written in the Python programming language (File S2).

### A.2 Derivation of the deterministic dynamics

The derivation of the deterministic dynamics is identical to Santer and Uecker (2020). We recapitulate it here such that the article is self-contained.

To obtain the deterministic dynamics, we look at all events that alter the number of cells *N_j_* of a distinct type *j*. The following events can alter *N_j_*: 1) Cell division of *j*-type cells, which occurs at rate *N_j_λ_j_* and reduces *N_j_* by 1. 2) When cells of any type *i* divide, they may produce *m* = 1 or *m* =2 daughter cells of type *j*, which increases *N_j_* by *m*, with probability

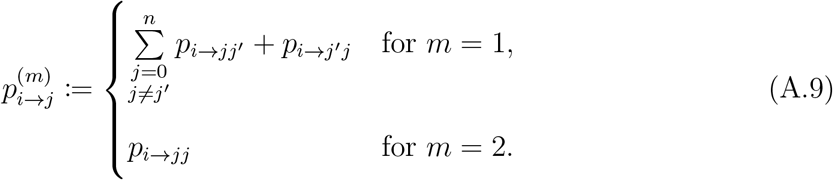

3) Replacement of a randomly chosen cell of type *j*, which reduces *N_j_* by 1. All combinations of these three events with the corresponding rates are listed in Table 1. Putting all those terms together, we obtain

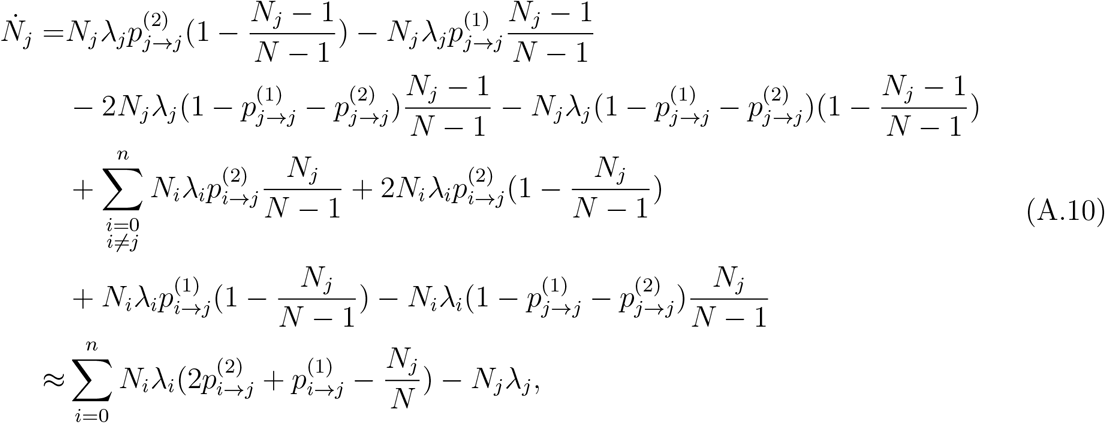

where we have used *N* – 1 ≈ *N* and *N_j_* – 1 ≈ *N_j_* (Santer and Uecker, 2020). The deterministic dynamics of the system is obtained by simultaneously integrating all *n* +1 equations for the cell type frequencies 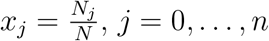:

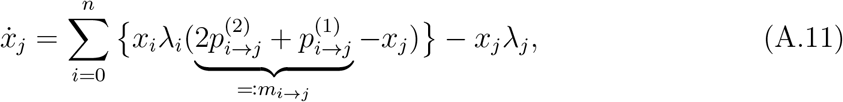

where we introduced *m*_*i→j*_ as the expected number of *j*-type cells produced at division of an i-type cell used below.

**Table 1:** Events that involve cells of type *j*. A parental cell of type *i* produces two daughter cells of type {*j, j*’} replacing the parental cell and one randomly chosen cell of type *l*. Rates of the events are obtained by the product of cell division rates and the probability that a *j*-type cell is replaced. *j, = j* denotes that one of the two daughter cells is of type *j* and the other is not of type *j* regardless of the order.

| Parental<br>cell type $i$ | Daughter<br>cell types | Cell type of<br>replaced cell | Change<br>in $N_j$ | Rate of<br>reaction |
| --- | --- | --- | --- | --- |
| $j$ | $j, j$ | $j$ | 0 | — |
| $j$ | $j, j$ | $\neq j$ | +1 | $N_j \lambda_j p_{j \rightarrow j}^{(2)} \left(1 - \frac{N_j - 1}{N - 1}\right)$ |
| $j$ | $j, \neq j$ | $j$ | -1 | $N_j \lambda_j p_{j \rightarrow j}^{(1)} \frac{N_j - 1}{N - 1}$ |
| $j$ | $j, \neq j$ | $\neq j$ | 0 | — |
| $j$ | $\neq j, \neq j$ | $j$ | -2 | $N_j \lambda_j (1 - p_{j \rightarrow j}^{(1)} - p_{j \rightarrow j}^{(2)}) \frac{N_j - 1}{N - 1}$ |
| $j$ | $\neq j, \neq j$ | $\neq j$ | -1 | $N_j \lambda_j (1 - p_{j \rightarrow j}^{(1)} - p_{j \rightarrow j}^{(2)}) \left(1 - \frac{N_j - 1}{N - 1}\right)$ |
| $\neq j$ | $j, j$ | $j$ | +1 | $N_i \lambda_i p_{i \rightarrow j}^{(2)} \frac{N_j}{N - 1}$ |
| $\neq j$ | $j, j$ | $\neq j$ | +2 | $N_i \lambda_i p_{i \rightarrow j}^{(2)} \left(1 - \frac{N_j}{N - 1}\right)$ |
| $\neq j$ | $j, \neq j$ | $j$ | 0 | — |
| $\neq j$ | $j, \neq j$ | $\neq j$ | +1 | $N_i \lambda_i p_{i \rightarrow j}^{(1)} \left(1 - \frac{N_j}{N - 1}\right)$ |
| $\neq j$ | $\neq j, \neq j$ | $j$ | -1 | $N_i \lambda_i (1 - p_{i \rightarrow j}^{(1)} - p_{i \rightarrow j}^{(2)}) \frac{N_j}{N - 1}$ |
| $\neq j$ | $\neq j, \neq j$ | $\neq j$ | 0 | — |

### A.3 Mathematical derivation for the conditions under which a heterozygosity window exists under random segregation

We here derive the criteria (1) and (2) for observing a heterozygosity window given that the initial frequency of mutant cells *f* is small. By definition, a heterozygosity window occurs if there are still heterozygous cells present at the time of fixation *t*_phen_ at the phenotype level. This may simply happen if the heterozygous cells that were present at time *t* = 0 have not fully decayed yet. This leads to a very small window though. A more relevant heterozygosity window occurs, if the heterozygotes initially increase in frequency during the fixation process (Figure 2). This observation builds the basis of our approximation.

For the frequency of heterozygous cells, i.e., 0 < *j* < *n*, we derive from Eq. (A.11)

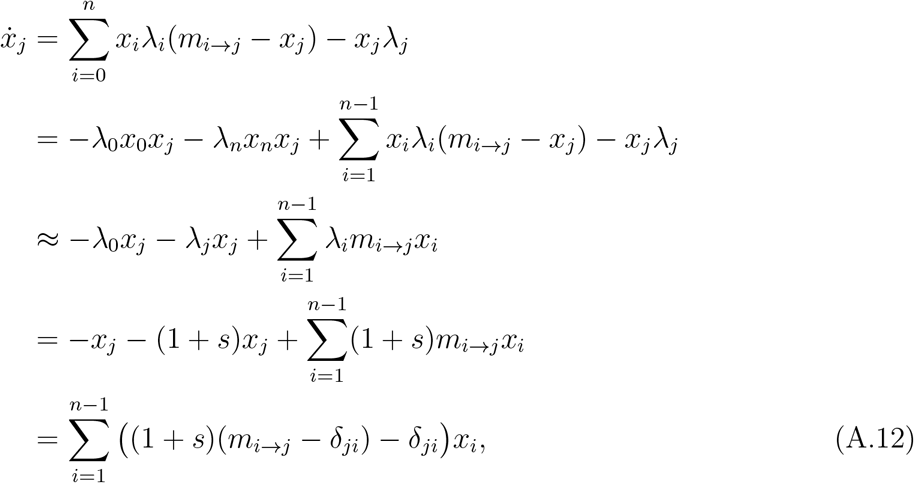

where *δ_ji_* denotes Kronecker’s Delta. Here, we have neglected quadratic terms of mutant cell frequencies *x_i_x_j_*, where *i* > 0 and *j* > 0, as mutant cell frequencies are low at early time points in the fixation process.

We choose the initial mutant frequency *f* sufficiently low so that the relative frequencies of heterozygote types equilibrate quickly compared to the time it takes for mutant cells to reach high frequencies in the population. The frequencies of heterozygote types then take approximately the form of the right eigenvector corresponding to the leading eigenvalue of the matrix (*m_i→j_* – *δ_ji_*)_*ji*∈{1,…,*n*_1}_ (see Eq. (A.12)). In the following, we denote by *x_i_* the frequencies of cells of type *i* at such an early time t assuming that the relative heterozygous frequencies are in equilibrium.

If replication is regular, we have 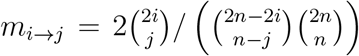 (see Eq. (A.4) and (A.9)). The dominant eigenvalue of the matrix (*m*_*i→j*_ – *δ_ij_*)_*ji*∈{1,…,*n*-1}_ for regular replication can be calculated explicitely as

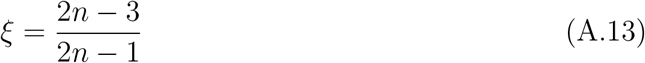

(Novick and Hoppensteadt, 1978, λ* in their Eq. (2) is the dominant eigenvalue of the matrix (*m_i→j_*/2)_*i,j*∈{1,..,*n*_1}_). Since (*x*_1_,…, *x*_*n*-1_) is the eigenvector corresponding to *ξ*, we obtain for the time derivative of the frequency of all heterozygous cells 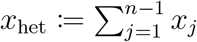:

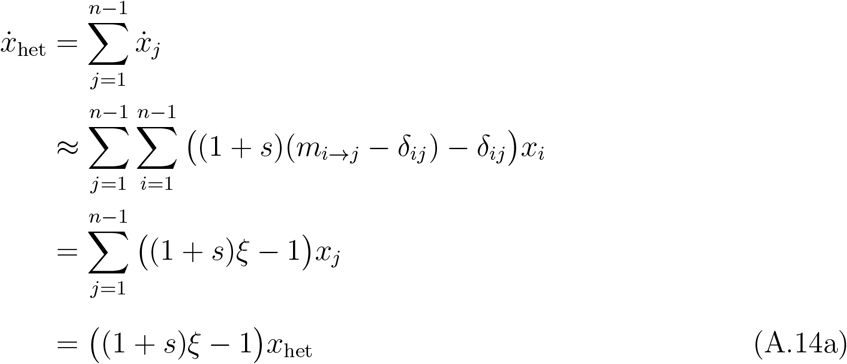

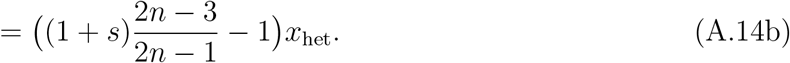

Thus, heterozygous cells increase in frequency early in the fixation process if

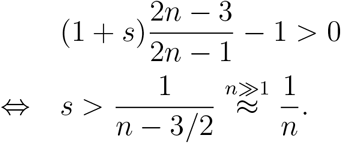

If there is no selection, i.e., *s* = 0, we have

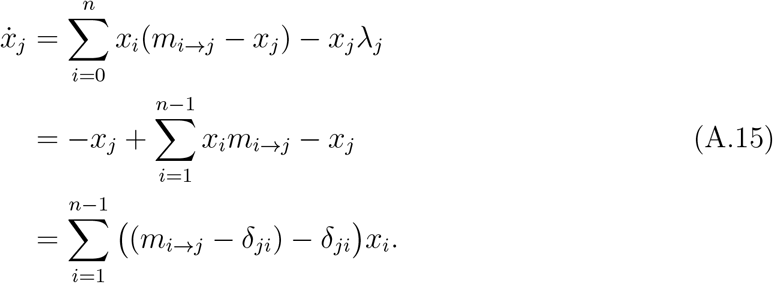

(The final expression is the same as in Eq. (A.12), but the derivation does not require any approximation if *s* = 0.) Analogous to Eq. (A.14b), we then obtain

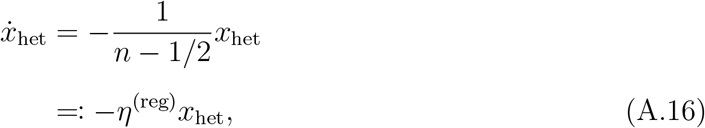

where *η*^(reg)^ can be interpreted as the heterozygote loss rate.

For random replication, the dominant eigenvalue of the matrix (*m*_i→j_ – *δ_ij_*)_*ji*∈{1,…,*n*-1}_ can be calculated as

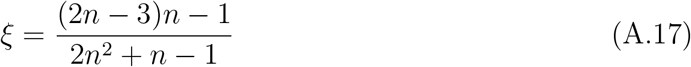

(Novick and Hoppensteadt, 1978). For this mode, we obtain in an analogous manner the criterion for the heterozygosity window

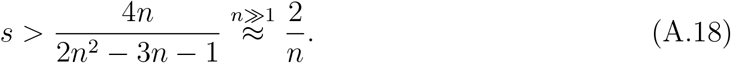

and the heterozygote loss rate

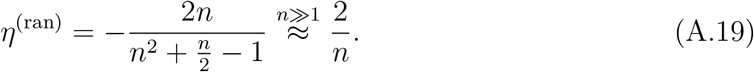

In a simpler model used by Rodriguez-Beltran et al. (2018), where heterozygous cells always contain an equal proportion of mutant and wild-type copies, the dynamics of heterozygous cells without selection and at low heterozygote frequencies can be described analogous to Eq. (A.12) by

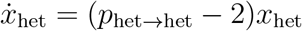

where *p*_het→het_ = 2(1 – 2^1-*n*^) denotes the expected number of heterozygous cells created at cell division of a heterozygote. Note that 2^1-*n*^ is the probability that two homozygous cells are formed at cell division of a heterozygous cell (Rodriguez-Beltran et al., 2018). This leads to

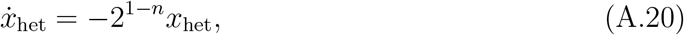

where 2^1-*n*^ is the heterozygote loss rate under this simpler model. Here, the heterozygote loss rate decreases exponentially with the replicon copy number *n*, whereas in our model, which considers different compositions of mutant and wild-type copies, the loss rate scales with 1/*n*.

### A.4 Cell-type frequencies at transformation-selection balance

In this section, we derive the population composition at transformation-selection balance. We assume that at every transformation event, one replicon copy in the recipient cell is replaced. Mutant copies are integrated at per cell rate *τ*. For a cell with *j* mutant replicon copies, the probability that a wild-type copy is replaced is given by 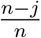. This changes its cell type *j* → *j* + 1. Mutant cells of type *j* > 0 have a reduced reproduction rate 1 – *σ* < 1, i.e., they are negatively selected. The transformation-selection balance reflects the state at which the input of the mutant variant through transformation and the selection outweigh each other. The time derivatives of the cell-type abundances 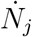 are given by

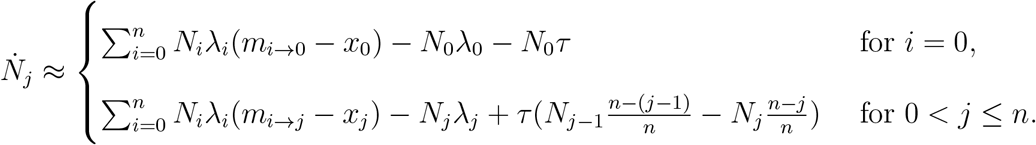

The time derivatives of the relative frequencies *x_i_* = *N_i_/N* are thus given by

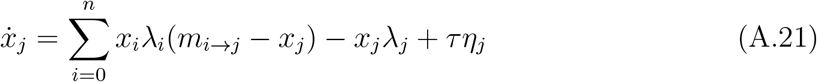

with 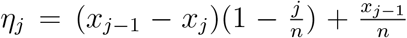 for *i* > 0 and *η*_0_ = – *x*_0_. Finding the equilibrium (*x*_0_,…, *x_n_*) where 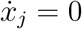 for all *j* was performed numerically by simultaneously integrating *x_j_* for all *j* until 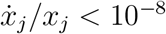.

## References

C. Bailey, M. Shoura, P. Mischel, and C. Swanton. Extrachromosomal DNA—relieving heredity constraints, accelerating tumour evolution. Annals of Oncology, 31(7):884–893, 2020.

J. A. Bogan, J. E. Grimwade, M. Thornton, P. Zhou, G. D. Denning, and C. E. Helmstetter. P1 and NR1 plasmid replication during the cell cycle of escherichia coli. Plasmid, 45(3):200–208, 2001.

S. Breuert, T. Allers, G. Spohn, and J. Soppa. Regulated polyploidy in halophilic archaea. PLoS ONE, 1(1):e92, 2006.

A. C. Brooks and L. C. Hwang. Reconstitutions of plasmid partition systems and their mechanisms. Plasmid, 91:37–41, 2017.

J. Cullum and P. Broda. Rate of segregation due to plasmid incompatibility. Genetical Research, 33(1):61–79, 1979.

F. Eggenberger and G. Pólya. Über die Statistik verketteter Vorgänge. ZAMM – Journal of Applied Mathematics and Mechanics / Zeitschrift für Angewandte Mathematik und Mechanik, 3(4):279–289, 1923.

K. Friehs. Plasmid copy number and plasmid stability. In T. Scheper, editor, New Trends and Developments in Biochemical Engineering, pages 47–82. Springer, Berlin, Heidelberg, 2004.

A. Garoña, N. F. Hülter, D. R. Picazo, and T. Dagan. Segregational drift constrains the evolutionary rate of prokaryotic plasmids. Molecular Biology and Evolution, 38(12):5610–5624, 2021.

D. T. Gillespie. A general method for numerically simulating the stochastic time evolution of coupled chemical reactions. Journal of Computational Physics, 22:403–434, 1976.

M. Griese, C. Lange, and J. Soppa. Ploidy in cyanobacteria. FEMS Microbiology Letters, 323(2):124–131, 2011.

F. Guerrini and M. S. Fox. Genetic heterozygosity in pneumococcal transformation. Pro-ceedings of the National Academy of Sciences, 59(2):429–436, 1968.

A. D. Halleran, E. Flores-Bautista, and R. M. Murray. Quantitative characterization of random partitioning in the evolution of plasmid-encoded traits. bioRxiv, 2019.

C. Hildenbrand, T. Stock, C. Lange, M. Rother, and J. Soppa. Genome copy numbers and gene conversion in methanogenic archaea. Journal of Bacteriology, 193(3):734–743, 2011.

B. Hu, G. Yang, W. Zhao, Y. Zhang, and J. Zhao. MreB is important for cell shape but not for chromosome segregation of the filamentous cyanobacterium Anabaena sp. PCC 7120. Molecular Microbiology, 63(6):1640–1652, 2007.

J. Ilhan, A. Kupczok, C. Woehle, T. Wein, N. F. Hülter, P. Rosenstiel, G. Landan, I. Mizrahi, and T. Dagan. Segregational drift and the interplay between plasmid copy number and evolvability. Molecular Biology and Evolution, 36(3):472–486, 2019.

D. Ionescu, M. Bizic-Ionescu, N. D. Maio, H. Cypionka, and H.-P. Grossart. Communitylike genome in single cells of the sulfur bacterium Achromatium oxaliferum. Nature Communications, 8(1):455, 2017.

K. Ishii, T. Hashimoto-Gotoh, and K. Matsubara. Random replication and random assortment model for plasmid incompatibility in bacteria. Plasmid, 1(4):435–445, 1978.

V. N. Iyer. Unstable genetic transformation in Bacillus subtilis and the mode of inheritance in unstable clones. Journal of Bacteriology, 90(2):495–503, 1965.

I. H. Jain, V. Vijayan, and E. K. O’Shea. Spatial ordering of chromosomes enhances the fidelity of chromosome partitioning in cyanobacteria. Proceedings of the National Academy of Sciences, 109(34):13638–13643, 2012. doi: 10.1073/pnas.1211144109.

M. Kimura. Some problems of stochastic processes in genetics. The Annals of Mathematical Statistics, 28(4):882–901, 1957.

M. Kimura and T. Ohta. The average number of generations until fixation of a mutant gene in a finite population. Genetics, 61(3):763–71, 1969.

C. Lawless, L. Greaves, A. K. Reeve, D. M. Turnbull, and A. E. Vincent. The rise and rise of mitochondrial DNA mutations. Open Biology, 10(5):200061, 2020.

H. Li. Random chromosome partitioning in the polyploid bacterium Thermus thermophilus HB27. G3: Genes, Genomes, Genetics, 9(4):g3.400086.2019, 2019.

R. N. Lightowlers, P. F. Chinnery, D. M. Turnbull, and N. Howell. Mammalian mitochondrial genetics: heredity, heteroplasmy and disease. Trends in Genetics, 13(11):450–455, 1997.

H. Mahmoud. Polya Urn Models. Chapman & Hall/CRC, 1 edition, 2008.

R. Maldonado, J. Jiménez, and J. Casadesús. Changes of ploidy during the *Azotobacter vinelandii* growth cycle. Journal of Bacteriology, 176(13):3911–3919, 1994.

S. Million-Weaver and M. Camps. Mechanisms of plasmid segregation: Have multicopy plasmids been overlooked? Plasmid, 75:27–36, 2014.

P. A. P. Moran. Random processes in genetics. Mathematical Proceedings of the Cambridge Philosophical Society, 54(1):60–71, 1958.

D. W. Morgens and A. R. Cavalcanti. Amitotic chromosome loss predicts distinct patterns of senescence and non-senescence in ciliates. Protist, 166(2):224–233, 2015.

M. L. Morse, E. M. Lederberg, and J. Lederberg. Transduction in Escherichia Coli K-12. Genetics, 41(1):142–56, 1956.

K. M. Münch, J. Müller, S. Wienecke, S. Bergmann, S. Heyber, R. Biedendieck, R. Münch, and D. Jahn. Polar Fixation of Plasmids during Recombinant Protein Production in Bacillus megaterium Results in Population Heterogeneity. Applied and Environmental Microbiology, 81(17):5976–5986, 2015. doi: 10.1128/aem.00807-15.

H. J. Nielsen, B. Youngren, F. G. Hansen, and S. Austin. Dynamics of Escherichia coli chromosome segregation during multifork replication. Journal of Bacteriology, 189(23):8660–8666, 2007.

K. Nordström. Plasmid R1—replication and its control. Plasmid, 55(1):1–26, 2006.

K. Nordsträom and S. Dasgupta. Copy-number control of the Escherichia coli chromosome: a plasmidologist’s view. EMBO reports, 7(5):484–489, 2006.

K. Nordström and K. Gerdes. Clustering versus random segregation of plasmids lacking a partitioning function: a plasmid paradox? Plasmid, 50(2):95–101, 2003.

R. P. Novick and F. Hoppensteadt. On plasmid incompatibility. Plasmid, 1(4):421–434, 1978.

R. Ohbayashi, A. Nakamachi, T. S. Hatakeyama, S. Watanabe, Y. Kanesaki, T. Chibazakura, H. Yoshikawa, S. Miyagishima, J. Soppa, and R. Losick. Coordination of polyploid chromosome replication with cell size and growth in a cyanobacterium. mBio, 10(2):e00510–19, 2019.

Y. Pichugin, W. Huang, and B. Werner. Stochastic dynamics of extra-chromosomal DNA. bioRxiv, 2019.

L. Pulakat, E. T. Efuet, and N. Gavini. Segregation pattern of kanamycin resistance marker in *Azotobacter vinelandii* did not show the constraints expected in a polyploid bacterium. FEMS Microbiology Letters, 160(2):247–252, 1998.

A. J. Ramsey and J. R. Mandel. When One Genome Is Not Enough: Organellar Heteroplasmy in Plants, pages 619–658. John Wiley & Sons, Ltd, 2019.

J. Rodríguez-Beltrán, J. DelaFuente, R. León-Sampedro, R. C. MacLean, and A. San Millan. Beyond horizontal gene transfer: the role of plasmids in bacterial evolution. Nature Reviews Microbiology, 19:347–359, 2021. doi: doi.org/10.1038/s41579-020-00497-1.

J. Rodriguez-Beltran, J. C. R. Hernandez-Beltran, J. DelaFuente, J. A. Escudero, A. Fuentes-Hernandez, R. C. MacLean, R. Peña-Miller, and A. S. Millan. Multicopy plasmids allow bacteria to escape from fitness trade-offs during evolutionary innovation. Nature Ecology & Evolution, 2(5):873–881, 2018.

R. Rownd. Replication of a bacterial episome under relaxed control. Journal of Molecular Biology, 44(3):387–402, 1969.

M. Santer and H. Uecker. Evolutionary rescue and drug resistance on multicopy plasmids. Genetics, 215(3):847–868, 2020.

D. Schneider, E. Fuhrmann, I. Scholz, W. R. Hess, and P. L. Graumann. Fluorescence staining of live cyanobacterial cells suggest non-stringent chromosome segregation and absence of a connection between cytoplasmic and thylakoid membranes. BMC Cell Biology, 8, 2007.

K. Skarstad, E. Boye, and H. Steen. Timing of initiation of chromosome replication in individual Escherichia coli cells. The EMBO Journal, 5(7):1711–1717, 1986.

J. Soppa. Non-equivalent genomes in polyploid prokaryotes. Nature Microbiology, 2021. doi: doi.org/10.1038/s41564-021-01034-3.

J. Soppa. Polyploidy and community structure. Nature Microbiology, 2(2):16261, 2017.

J. B. Stewart and P. F. Chinnery. The dynamics of mito chondrial DNA heteroplasmy: implications for human health and disease. Nature Reviews Genetics, 16(9):530–542, 2015.

M. H. Suh, L. Pulakat, and N. Gavini. Isolation and characterization of nif::kanamycin and nitrogen fixation proficient *Azotobacter vinelandii* strain, and its implication on the status of multiple chromosomes in azotobacter. Genetica, 110:101–107, 2000.

L. Sun, H. K. Alexander, B. Bogos, D. J. Kiviet, M. Ackermann, and S. Bonhoeffer. Effective polyploidy causes phenotypic delay and influences bacterial evolvability. PLOS Biology, 16(2):e2004644, 2018.

D. M. Tobiason and H. S. Seifert. Genomic content of *Neisseria species*. Journal of Bacteriology, 192(8):2160–2168, 2010.

X. Wang, H. B. Brandão, T. B. K. Le, M. T. Laub, and D. Z. Rudner. Bacillus subtilis SMC complexes juxtapose chromosome arms as they travel from origin to terminus. Science, 355(6324):524–527, 2017.

S. Watanabe. Cyanobacterial multi-copy chromosomes and their replication. Bioscience, Biotechnology, and Biochemistry, 84(7):1309–1321, 2020.

S. Watanabe, R. Ohbayashi, Y. Shiwa, A. Noda, Y. Kanesaki, T. Chibazakura, and H. Yoshikawa. Light-dependent and asynchronous replication of cyanobacterial multi-copy chromosomes. Molecular Microbiology, 83(4):856–865, 2012.

H.-Y. Wu, K. Lau, and L. F. Liu. Interlocking of plasmid DNAs due to lac repressoroperator interaction. Journal of Molecular Biology, 228(4):1104–1114, 1992.

