## Supplementary material for "Fixation dynamics of beneficial alleles in prokaryotic polyploid chromosomes and plasmids": File S1

### Fixation dynamics of beneficial alleles in prokaryotic polyploid chromosomes and plasmids
### Supplementary Information
#### File S1

Mario Santer[1], Anne Kupczok[2,3], Tal Dagan[2], and Hildegard Uecker[1]

[1]Research group Stochastic Evolutionary Dynamics, Department of Evolutionary Theory, Max Planck Institute for Evolutionary Biology, Plön, Germany

[2]Institute of General Microbiology, Kiel University, Kiel, Germany

[3]Bioinformatics group, Department of Plant Sciences, Wageningen University & Research, Wageningen, Netherlands

### S1: The mathematics and convergence of the heterozygosity window

In the following, we derive an analytical approximation for the solution of cell-type frequencies over time for the assumption that the initial frequency $f$ is very small, similar to Eq. (A.12) in section A.3. Here, we only consider the mode of random segregation. Later on we use this solution to proof the convergence of the heterozygosity window in the limit of the threshold $x_{\text{thr}} \to 1$ as discussed in the main text.

As in Appendix A.3, we choose the initial frequency $f$ sufficiently low such that the relative frequencies of heterozygotes $\chi_j := \frac{x_j}{x_1+\cdots+x_{n-1}}$ equilibrate at a timescale that is short relative to the time it takes until mutant cells take over the population. Hence, we assume in the following that $(x_1, \ldots, x_{n-1})^T$ is proportional to the eigenvector of $(m_{i\to j} - \delta_{ji})_{ji\in\{1,\ldots,n-1\}}$ corresponding to the dominant eigenvalue $\xi$, which depends on the copy number $n$ and on the mode of replication (see Eq. (A.13) and (A.17), $m_{i\to j}$ defined in (A.12) denotes the expected number of $j$-type cells produced at division of a $i$-type cell). Once equilibrated, the frequencies of the heterozygous cells relative to each other $\chi_j$ remain constant throughout the entire fixation process, which can formally be seen by the following calculation:

$$
\begin{aligned}
\frac{\mathrm{d}\frac{x_j}{x_\text{het}}}{\mathrm{d}t} &= \frac{\dot{x}_j}{x_\text{het}} - \frac{x_j}{x_\text{het}^2}\dot{x}_\text{het} \\
&= \frac{1}{x_\text{het}}\left\{\dot{x}_j - \frac{x_j}{x_\text{het}}x_\text{het}\right\} \\
&= \frac{1}{x_\text{het}}\left\{-x_0x_j - (1+s)x_nx_j + \sum_{i=1}^{n-1} x_i(1+s)(m_{i\to j} - x_j) - x_j(1+s) - \frac{x_j}{x_\text{het}}\sum_{k=1}^{n-1}\dot{x}_k\right\} \\
&= \frac{1}{x_\text{het}}\Bigg\{ - x_0x_j - (1+s)x_nx_j + \sum_{i=1}^{n-1} x_i(1+s)m_{i\to j} - (1+s)x_jx_\text{het} - x_j(1+s) \\
&\qquad - \frac{x_j}{x_\text{het}}\sum_{k=1}^{n-1}\left(-x_0x_k - (1+s)x_nx_k\sum_{i=1}^{n-1} x_i(1+s)(m_{i\to k} - x_k) - x_k(1+s)\right)\Bigg\} \\
&= \frac{1}{x_\text{het}}\Bigg\{ - x_0x_j - (1+s)x_j(1-x_0) + \sum_{i=1}^{n-1}(1+s)x_i(m_{i\to j} - \delta_{ij}) \\
&\qquad - \frac{x_j}{x_\text{het}}\sum_{k=1}^{n-1}\left(-x_0x_k - (1+s)x_k(1-x_0) + \sum_{i=1}^{n-1} x_i(1+s)(m_{i\to k} - \delta_{ik})\right)\Bigg\} \\
&= \frac{1}{x_\text{het}}\Bigg\{ - x_0x_j - (1+s)x_j(1-x_0) + \sum_{i=1}^{n-1}(1+s)x_i(m_{i\to j} - \delta_{ij}) \\
&\qquad - \frac{x_j}{x_\text{het}}\left(-x_0x_\text{het} - (1+s)x_\text{het}(1-x_0) + \sum_{k=1}^{n-1}\sum_{i=1}^{n-1} x_i(1+s)(m_{i\to k} - \delta_{ik})\right)\Bigg\} \\
&= \frac{1}{x_\text{het}}\left\{\sum_{i=1}^{n-1}(1+s)x_i(m_{i\to j} - \delta_{ij}) - \frac{x_j}{x_\text{het}}\sum_{k=1}^{n-1}\sum_{i=1}^{n-1} x_i(1+s)(m_{i\to k} - \delta_{ik})\right\} \\
&= (1+s)\left\{\sum_{i=1}^{n-1}\frac{x_i}{x_\text{het}}(m_{i\to j} - \delta_{ij}) - \frac{x_j}{x_\text{het}}\sum_{k=1}^{n-1}\sum_{i=1}^{n-1}\frac{x_i}{x_\text{het}}(m_{i\to k} - \delta_{ik})\right\} \\
&= (1+s)\left\{\xi\frac{x_j}{x_\text{het}} - \frac{x_j}{x_\text{het}}\sum_{k=1}^{n-1}\xi\frac{x_k}{x_\text{het}}\right\} \\
&= (1+s)\left\{\xi\frac{x_j}{x_\text{het}} - \frac{x_j}{x_\text{het}}\xi\right\} \\
&= 0. \qquad \text{(S1.1)}
\end{aligned}
$$

The evolution of the population through time in our model can then be described by a system of three ordinary differential equations for the frequency of the wild type $x_0$, the sum of frequencies of all heterozygous types $x_{\text{het}} = x_1 + \cdots + x_{n-1}$, and frequency of the homozygous mutant type $x_n$. From equation (A.11), we obtain for the time-derivative of the homozygous mutant type $i = n$

$$\begin{aligned}
\dot{x}_n &= \sum_{i=0}^{n} \{x_i \lambda_i (m_{i\to n} - x_n)\} - x_i \lambda_n \\
&= x_0(m_{0\to n} - x_n) + \sum_{i=1}^{n-1} x_i(1+s)(m_{i\to n} - x_n) + x_n(1+s)(m_{n\to n} - x_n) - x_n(1+s) \\
&= -x_0 x_n + (1+s)\left(\sum_{i=1}^{n-1} m_{i\to n}\chi_i x_{\text{het}} - x_n \sum_{i=1}^{n-1} x_i\right) + x_n(1+s)(2 - x_n) - x_n(1+s) \\
&= -x_0 x_n + (1+s)\kappa x_{\text{het}} - (1+s)x_{\text{het}} x_n - (1+s)x_n^2 + (1+s)x_n \\
&= -(1 - x_{\text{het}} - x_n) + (1+s)\kappa x_{\text{het}} - (1+s)x_{\text{het}} x_n - (1+s)x_n^2 + (1+s)x_n \\
&= -s x_n^2 - s x_{\text{het}} x_n + (1+s)\kappa x_{\text{het}} + s x_n,
\end{aligned}$$

where we used $m_{0\to n} = 0$ and $m_{n\to n} = 2$, and defined $\kappa := \sum_{i=1}^{n-1} m_{i\to n} \frac{x_i}{x_{\text{het}}}$. It holds

$$\begin{aligned}
\kappa &= \sum_{i=1}^{n-1} (m_{i\to n} - \delta_{in})\chi_i \\
&= \sum_{j=0}^{n}\sum_{i=1}^{n-1} (m_{i\to j} - \delta_{ij})\chi_i - \sum_{j=0}^{n-1}\sum_{i=1}^{n-1} (m_{i\to j} - \delta_{ij})\chi_i \\
&= 1 - \sum_{i=1}^{n-1} (m_{i\to 0} - \delta_{ij})\chi_i - \sum_{j=1}^{n-1}\sum_{i=1}^{n-1} (m_{i\to j} - \delta_{ij})\chi_i \\
&= 1 - \sum_{i=1}^{n-1} (m_{i\to n} - \delta_{ij})\chi_i - \xi \\
&= 1 - \kappa - \xi,
\end{aligned}$$

where we have used $\sum_{i=1}^{n-1}\sum_{j=0}^{n} \left(m_{i\to j} - \delta_{ij}\right)\chi_i = 1$ (every cell has two daughter cells) and the symmetry $m_{i\to j} = m_{n-i\to n-j}$. We therefore obtain

$$\kappa = \frac{1}{2}(1-\xi). \tag{S1.2}$$

For the sum of heterozygous cells, we obtain

$$\begin{aligned}
\dot{x}_{\text{het}} &= \sum_{i=1}^{n-1} \dot{x}_i \\
&= \sum_{i=1}^{n-1} \left( \sum_{k=0}^{n} x_k \lambda_k (m_{k\to i} - x_i) - x_i \lambda_i \right) \\
&= \sum_{i=1}^{n-1} \left( x_0 (m_{0\to i} x_i) + \sum_{k=1}^{n-1} x_k (1+s)(m_{k\to i} - x_i) + x_n (1+s)(m_{n\to i} - x_i) - x_i(1+s) \right) \\
&= \sum_{i=1}^{n-1} \left( -x_0 x_i + (1+s) \left( \sum_{k=1}^{n-1} m_{k\to i} \chi_k x_{\text{het}} - x_i \sum_{k=1}^{n-1} x_k \right) - x_n (1+s) x_i - x_i (1+s) \right) \\
&= \sum_{i=1}^{n-1} \left( -x_0 x_{\text{het}} + (1+s)(\xi+1)\chi_i x_{\text{het}} - (1+s) x_i x_{\text{het}} - x_n (1+s) x_i - x_i(1+s) \right) \\
&= -x_0 x_{\text{het}} + (1+s)(\xi+1) x_{\text{het}} - (1+s) x_{\text{het}}^2 - x_{\text{het}} (1+s) x_n - x_{\text{het}}(1+s) \\
&= -(1 - x_{\text{het}} - x_n) x_{\text{het}} + (1+s)(\xi+1) x_{\text{het}} + (1+s) x_{\text{het}}^2 - x_n (1+s) x_{\text{het}} - x_{\text{het}}(1+s) \\
&= -s x_{\text{het}}^2 - s x_n x_{\text{het}} + \left( (1+s)\xi - 1 \right) x_{\text{het}}.
\end{aligned}$$

To make progress, it is easier to switch variables and to consider the total frequency of mutant cells $x_{\text{mut}} \coloneqq x_{\text{het}} + x_{\text{n}}$ and the relative fraction of heterozygous cells among all mutant cells $x_{\mathrm{hf}} := \frac{x_{\mathrm{het}}}{x_{\mathrm{mut}}}$ instead of $x_{\mathrm{het}}$ and $x_n$. The time-derivative of $x_{\mathrm{mut}}$ is given by

$$
\begin{aligned}
\frac{\mathrm{d}x_{\mathrm{mut}}}{\mathrm{d}t} &= \dot{x}_{\mathrm{het}} + \dot{x}_n \\
&= -sx_{\mathrm{het}}^2 - sx_n^2 - 2sx_n x_{\mathrm{het}} + \left((1+s)(\xi+\kappa)-1\right) x_{\mathrm{het}} + sx_n \\
&= -s(x_{\mathrm{het}} + x_n)^2 + \left((1+s)(\xi+\kappa)-1\right) x_{\mathrm{het}} \\
&= -sx_{\mathrm{mut}}^2 + \left((1+s)(\xi+\kappa)-1\right) x_{\mathrm{mut}} x_{\mathrm{hf}} + sx_{\mathrm{mut}}(1-x_{\mathrm{hf}}) && \text{(S1.3a)} \\
&= -sx_{\mathrm{mut}}^2 + sx_{\mathrm{mut}} + \underbrace{(1+s)(\xi+\kappa-1)}_{=:a} x_{\mathrm{mut}} x_{\mathrm{hf}} && \text{(S1.3b)}
\end{aligned}
$$

For the time-derivative of $x_{\mathrm{hf}}$, we obtain

$$
\begin{aligned}
\dot{x}_{\mathrm{hf}} &= \frac{\mathrm{d}}{\mathrm{d}t}\left(\frac{x_{\mathrm{het}}}{x_{\mathrm{mut}}}\right) \\
&= \frac{\dot{x}_{\mathrm{het}}}{x_{\mathrm{mut}}} - \frac{x_{\mathrm{hf}}}{x_{\mathrm{mut}}}\dot{x}_{\mathrm{mut}} \\
&= \frac{-sx_{\mathrm{het}}^2 - sx_n x_{\mathrm{het}} + \left((1+s)\xi - 1\right) x_{\mathrm{het}}}{x_{\mathrm{mut}}} \\
&\quad - \frac{x_{\mathrm{hf}}}{x_{\mathrm{mut}}}\left(-sx_{\mathrm{mut}}^2 + \left((1+s)(\xi+\kappa)-1\right) x_{\mathrm{mut}} x_{\mathrm{hf}} + sx_{\mathrm{mut}}(1-x_{\mathrm{hf}})\right) \\
&= -sx_{\mathrm{het}} x_{\mathrm{hf}} - sx_n x_{\mathrm{hf}} + \left((1+s)\xi - 1\right) x_{\mathrm{hf}} \\
&\quad + sx_{\mathrm{hf}} x_{\mathrm{mut}} - \left((1+s)(\xi+\kappa)-1\right) x_{\mathrm{hf}}^2 - s(1-x_{\mathrm{hf}})x_{\mathrm{hf}} \\
&= -sx_{\mathrm{mut}} x_{\mathrm{hf}} + \left((1+s)\xi - 1\right) x_{\mathrm{hf}} \\
&\quad + sx_{\mathrm{hf}} x_{\mathrm{mut}} - \left((1+s)(\xi+\kappa)-1\right) x_{\mathrm{hf}}^2 - s(1-x_{\mathrm{hf}})x_{\mathrm{hf}} \\
&= \left((1+s)\xi - 1 - s\right)x_{\mathrm{hf}} + \left(s + 1 - (1+s)(\xi+\kappa)\right) x_{\mathrm{hf}}^2 \\
&= -\underbrace{(1+s)(\xi+\kappa-1)}_{=a} x_{\mathrm{hf}}^2 + \underbrace{(1+s)(\xi-1)}_{=:b} x_{\mathrm{hf}},
\end{aligned}
$$

where we have used Eq. (S1.3a). The general solution to this Bernoulli differential equation

(Zeidler, 2013) is

$$x_{\mathrm{hf}}(t) = \frac{be^{bC+bt}}{ae^{bC+bt}+1}. \tag{S1.4}$$

From the initial condition $x_1 = f, x_2 = 0, \ldots, x_n = 0$, such that $x_{\mathrm{hf}} = \frac{x_{\mathrm{het}}}{x_{\mathrm{mut}}} = 1$, we get $C = \frac{\log\left(\frac{1}{b-a}\right)}{b}$. Substituting $C$ into Eq. (S1.4) yields

$$\begin{aligned} x_{\mathrm{hf}}(t) &= \frac{b(\frac{1}{b-a})e^{bt}}{a(\frac{1}{b-a})e^{bt}+1} \\ &= \frac{be^{bt}}{ae^{bt}+b-a} \qquad \text{(S1.5a)} \\ &= \frac{(1+s)(\xi-1)e^{(1+s)(\xi-1)t}}{(1+s)(1-\xi-\kappa)e^{(1+s)(\xi-1)t}-(1+s)\kappa} \\ &= \frac{(\xi-1)e^{(1+s)(\xi-1)t}}{(1-\xi-\kappa)e^{(1+s)(\xi-1)t}-\kappa}. \qquad \text{(S1.5b)} \end{aligned}$$

Inserting Eq. (S1.5a) into (S1.3b) gives

$$\dot{x}_{\mathrm{mut}} = -sx_{\mathrm{mut}}^2 + sx_{\mathrm{mut}} + ax_{\mathrm{mut}}\frac{be^{bt}}{ae^{bt}+b-a}. \tag{S1.6}$$

For the initial condition $x_{\mathrm{mut}}(0) = f$, the solution (obtained with Mathematica Version 12.0.0.0 (Wolfram Research, Inc.)) is given by

$$\begin{aligned} x_{\mathrm{mut}}(t) &= \frac{f(b+s)e^{st}\left(a\left(e^{bt}-1\right)+b\right)}{fe^{st}\left(ase^{bt}-(a-b)(b+s)\right)+b(af+b(-f)+b-fs+s)} \\ &= \frac{f(\xi+\xi s-1)e^{st}\left(\kappa-(\kappa+\xi-1)e^{(\xi-1)(s+1)t}\right)}{\begin{aligned}&f\kappa(\xi-1)\left(e^{st}-1\right) \\ &\quad + fs\left(\kappa+\xi+\kappa\xi\left(e^{st}-1\right)-(\kappa+\xi-1)e^{t(\xi+\xi s-1)}-1\right)-(\xi-1)(\xi+\xi s-1)\end{aligned}}. \end{aligned} \tag{S1.7}$$

In the next step, we derive an expression for the size of the heterozygosity window in the limit $x_{\mathrm{thr}} \to 1$, where $x_{\mathrm{thr}}$ denotes the threshold for fixation, and show that it is independent of the initial frequency $f$. The limit $x_{\mathrm{thr}} \to 1$ implies that $t_{\mathrm{fix}}$ and $t_{\mathrm{phen}}$ both tend to infinity, and we thus study the behavior of the system for large times (formally $t \to \infty$).

For the time to fixation at the phenotype level $t_{\mathrm{phen}}$, it holds that $x_0(t_{\mathrm{phen}}) = 1 - x_{\mathrm{mut}}(t_{\mathrm{phen}}) = 1 - x_{\mathrm{thr}}$. For the time to fixation at the genotype level $t_{\mathrm{fix}}$ (fixation of homozygous mutant cells), it analogously holds $x_0(t_{\mathrm{fix}}) + x_{\mathrm{het}}(t_{\mathrm{fix}}) = 1 - x_{\mathrm{n}}(t_{\mathrm{fix}}) = 1 - x_{\mathrm{thr}}$, where we have $x_{\mathrm{wt}} \coloneqq x_0 + x_{\mathrm{het}}$ as the frequency of cells that carry wild-type replicon copies. Combining the latter equations, we get

$$1 - x_{\mathrm{thr}} = x_0(t_{\mathrm{phen}}) = x_{\mathrm{wt}}(t_{\mathrm{fix}}). \tag{S1.8}$$

If the strength of selection is large compared to the inverse copy number, so that we expect a heterozygosity window (cf. the threshold of Eq. (1)), the decay rate of the frequency of wild-type carrying cells $x_{\mathrm{wt}}(t)$ is approximately given by

$$\frac{1}{t}\log x_{\mathrm{wt}}(t) = \frac{1}{t}\log\left(1 - x_{\mathrm{n}}(t)\right) = \frac{1}{t}\log\left(1 - x_{\mathrm{mut}}(t)(1 - x_{\mathrm{hf}}(t))\right) \overset{t\to\infty}{\longrightarrow} (1+s)(\xi - 1), \tag{S1.9}$$

where we obtained the limit using Mathematica as above (see File S2). Mathematica states the condition $\xi > \frac{2+s}{2(1+s)}$ as a condition for this limit, which is is equivalent to $s > \frac{2}{n-5/2} \approx \frac{2}{n}$ for regular replication and $s > (8n)/(-1 - 7n + 2n^2) \approx \frac{4}{n}$ for random replication. This is more stringent than the condition for the existence of a heterozygosity window (Eq. (1)), which is $s \gtrsim \frac{1}{n}$ (regular replication, Eq. (1)) or $s \gtrsim \frac{2}{n}$ (random replication, Eq. (2)). Numerical results for relevant cases of $n$ and $s$, however, show that Eq. (S1.9) also holds true if $\frac{1}{n} \lesssim s \lesssim \frac{2}{n}$ (see File S2). Thus, for large times, we have

$$x_{\mathrm{wt}}(t) \propto e^{(1+s)(\xi-1)t}.$$

Consequently, we have

$$x_{\mathrm{wt}}(t_{\mathrm{fix}}) = x_{\mathrm{wt}}(t_{\mathrm{phen}})e^{(1+s)(\xi-1)\Delta t}, \tag{S1.10}$$

where we used the definition of the heterozygosity window $\Delta t = t_{\mathrm{fix}} - t_{\mathrm{phen}}$. Moreover, using again Mathematica, we obtain for the limit

$$\frac{x_{\mathrm{wt}}(t)}{x_0(t)} \overset{t\to\infty}{\longrightarrow} \frac{\kappa + (\kappa-1)s}{(1+s)(\kappa+\xi-1)} =: r, \tag{S1.11}$$

which is independent of $f$. Inserting Eq. (S1.10) and (S1.11) into Eq. (S1.8) gives for the limit of $t \to \infty$ where $x_{\mathrm{thr}} \to 1$

$$\begin{aligned}
& x_{\mathrm{wt}}(t_{\mathrm{phen}})e^{(1+s)(\xi-1)\Delta t} = x_{\mathrm{wt}}(t_{\mathrm{fix}}) = x_0(t_{\mathrm{phen}}) = \frac{x_{\mathrm{wt}}(t_{\mathrm{phen}})}{r} \\
\Leftrightarrow \quad & e^{(1+s)(\xi-1)\Delta t} = \frac{1}{r} \\
\Leftrightarrow \quad & \Delta t = \frac{\log\frac{1}{r}}{(1+s)(\xi-1)} = \frac{\log\left(\frac{(1+s)(\kappa+\xi-1)}{\kappa+(\kappa-1)s}\right)}{(1+s)(\xi-1)},
\end{aligned} \tag{S1.12}$$

which is independent of the initial frequency $f$.

For the mode of regular replication, we obtain for the heterozygosity window by insertion of the corresponding expressions for $\xi$ and $\kappa$ (Eqs. (A.13) and (S1.2))

$$\Delta t = \frac{2n-1}{2(1+s)} \ln\left(\frac{2(n-1)s-1}{s+1}\right) \tag{S1.13a}$$

$$\approx \frac{n}{1+s} \ln\left(\frac{2ns}{1+s}\right). \tag{S1.13b}$$

Under the mode of random replication, we obtain analogously using Eqs. (A.17) and (S1.2)

$$\Delta t = \frac{2n^2 + n - 1}{4n(1+s)} \ln\left(\frac{n(2ns - s - 2) - s}{2n(1+s)}\right) \tag{S1.14a}$$

$$\approx \frac{n/2}{1+s} \ln\left(\frac{ns}{1+s}\right). \tag{S1.14b}$$

### Supplementary figures

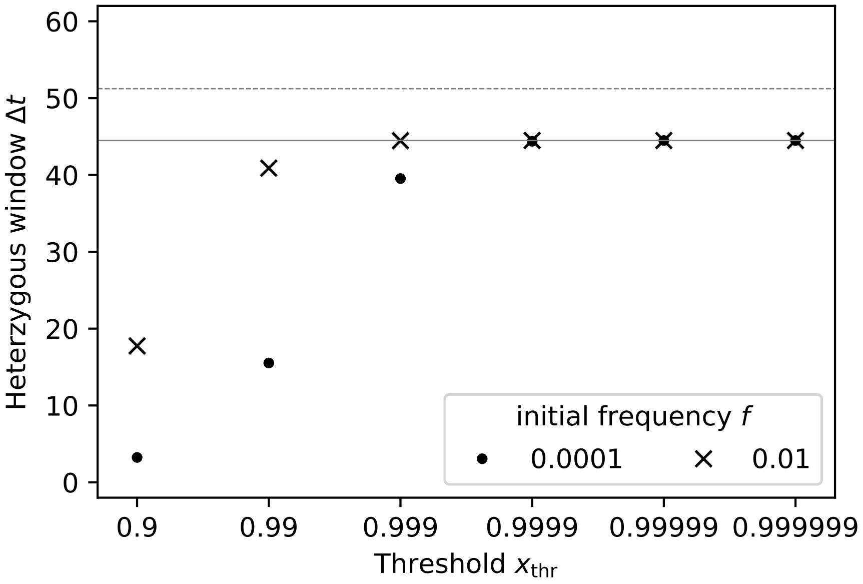


Figure S1: Influence of the fixation threshold $x_{\text{thr}}$ on the length of the heterozygosity window $\Delta t$. In the main text, a frequency of 99% mutant cells and of 99% homozygous mutant cells is the proxy for determining the fixation times $t_{\text{phen}}$ and $t_{\text{fix}}$ respectively. The plot shows the heterozygosity window $\Delta t = t_{\text{fix}} - t_{\text{phen}}$ for various thresholds $x_{\text{thr}}$ (dots) and the analytical approximations (solid and dashed lines, showing Eqs. (S1.13a) and (S1.13b), respectively). For smaller initial frequencies of mutant cells $f$ (see legend), the heterozygosity window converges later, i.e., for thresholds $x_{\text{thr}}$ closer to 1. Parameters: replicon copy number $n = 32$, strength of selection $s = 0.1$.

A

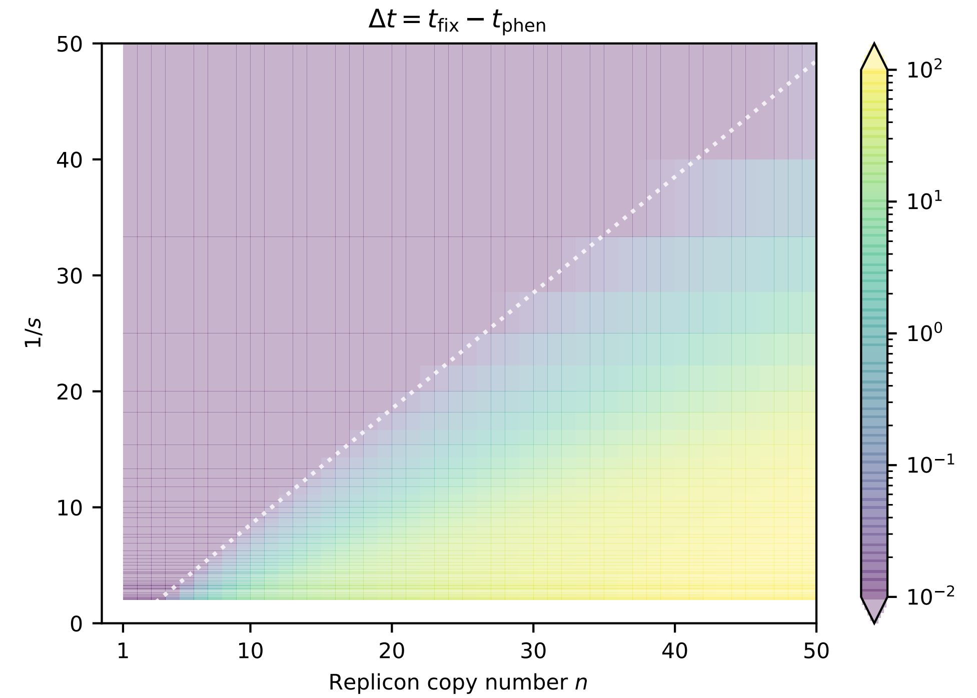


B

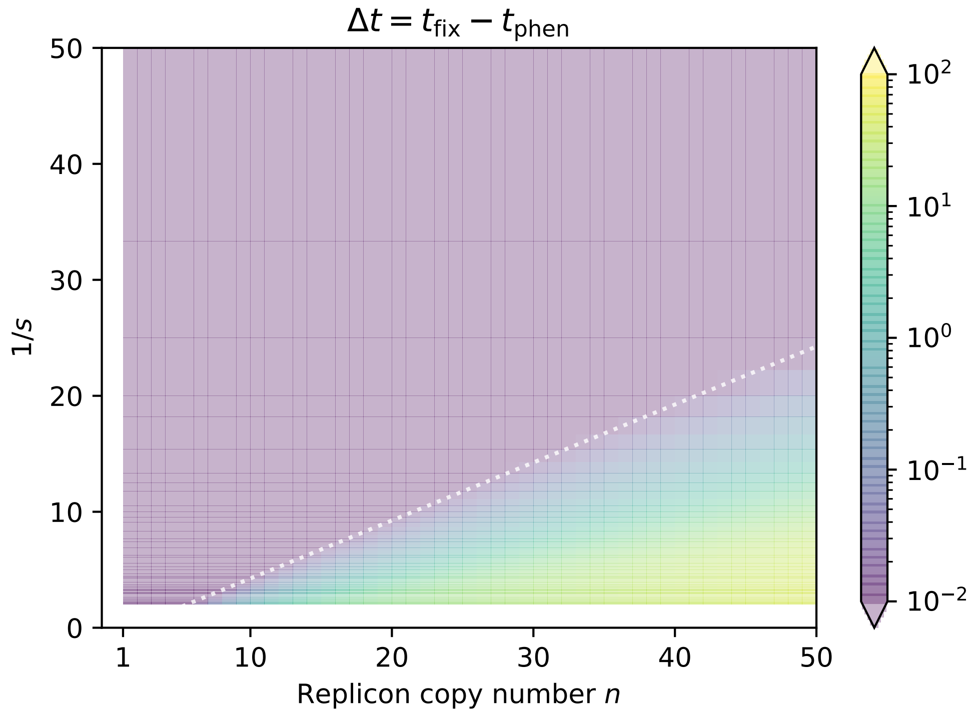


Figure S2: The heterozygosity window $\Delta t$ for various combinations of the replicon copy number $n$ and the inverse of the selective advantage $1/s$ for the mode of random segregation with (A) regular replication and (B) random replication. The initial frequency of mutant cells with one mutant replicon copy is $f = 0.01$. The dotted lines denote the threshold above which a heterozygosity windows arises given by Eq. (1) and (2) respectively.

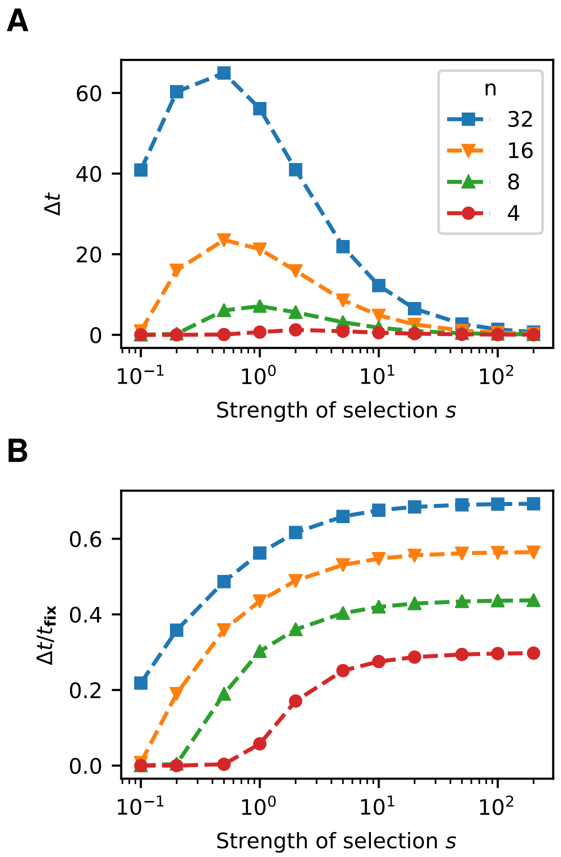


Figure S3: The heterozygosity window as a function of the strength of selection $s$ ($x$-axis) for various replicon copy numbers $n$ (lines). Panel A shows the absolute size $\Delta t$ and Panel B the size relative to the phenotypic fixation time $\Delta t/t_{\text{phen}}$. Both the absolute and the relative sizes increase with $n$. The relative size of the heterozygosity window (Panel B) also monotonically increases with the strength of selection $s$. The absolute size (Panel A) has a maximum as a function of $s$, since the fixation times become shorter with increasing $s$, which ulimately also affects the size of the window.

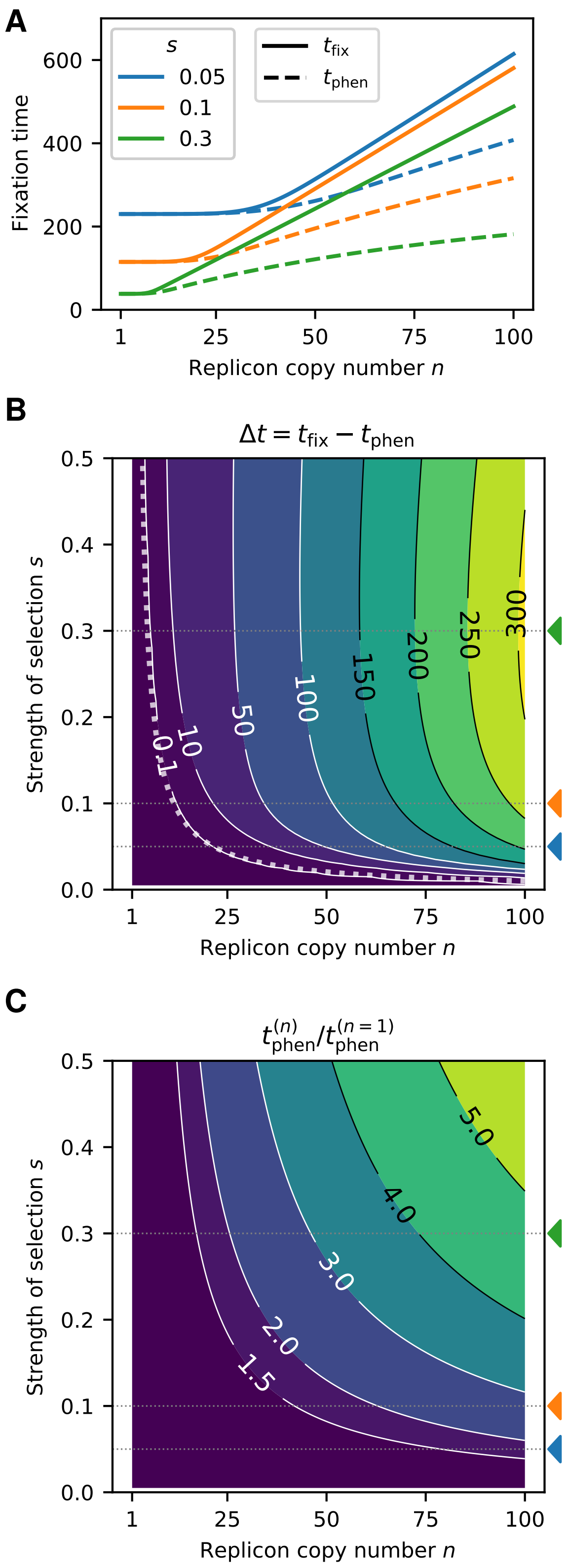


Figure S4: Influence of the replicon copy number $n$ and the strength of selection $s$ on the fixation times and the heterozygosity window for constant initial frequencies of mutant replicon copies $f_{rep} = f/n = 0.001$. The initial frequency of mutant cells with one mutant replicon copy is set to $f = f_{rep}n$. The panels are analogous to those of Figure 3, where $f$ rather than $f_{rep}$ is kept constant. (A) Fixation times as a function of the replicon copy number for several selection coefficients $s = 0.05$ (blue), 0.1 (orange), 0.3 (green) (B) Contour plot of the heterozygosity window for various replicon copy numbers $n$ and selection coefficients $s$. (C) Time of fixation at the phenotype level $t_{phen}$ relative to $n = 1$.

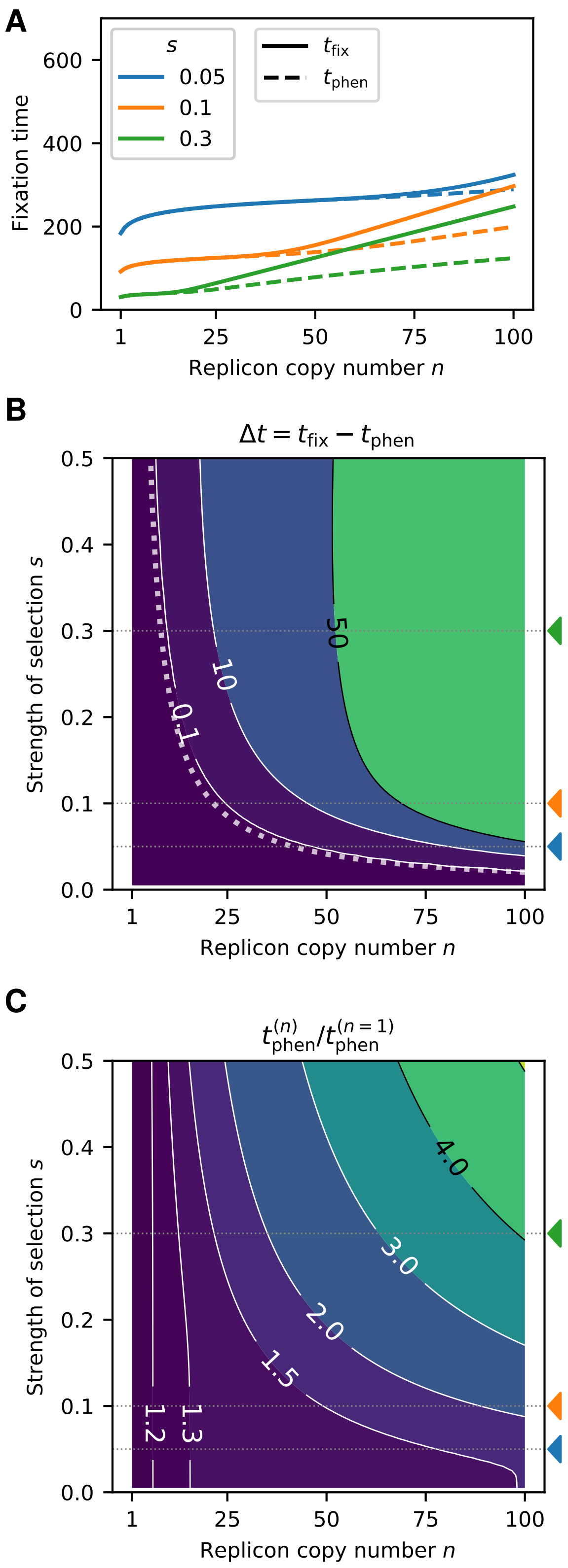


Figure S5: Influence of the replicon copy number $n$ and the strength of selection $s$ on the fixation times and the heterozygosity window for a replicon subject to random replication and random segregation. This figure is analogous to Figure 3, considering random replication instead of regular replication of replicon copies. The dotted line in (C) shows the threshold for $s$ at which the heterozygosity window start to occur (criterion (2)).

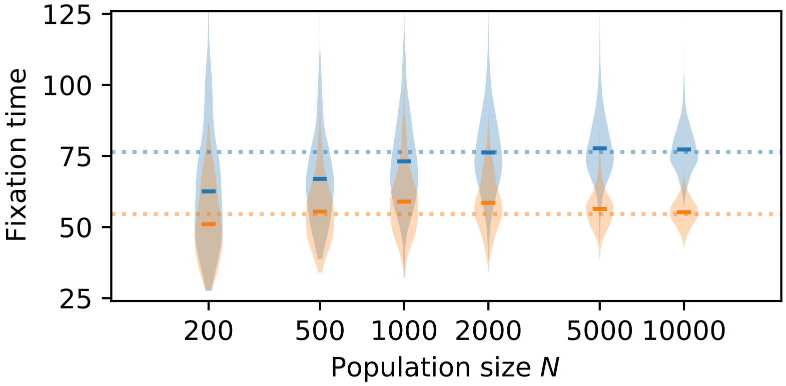


Figure S6: Fixation times of mutant cells $t_{\mathrm{phen}}$ (orange) and of homozygous mutant cells $t_{\mathrm{fix}}$ (blue) for various population sizes $N$. Violin plots show the distribution from $10^3$ stochastic simulations. Horizontal lines within the violin plots indicate the mean fixation times. The dashed horizontal lines show the deterministic results reflecting an infinite population. Parameters: $n = 16$, $s = 0.3$, $f = 0.01$.

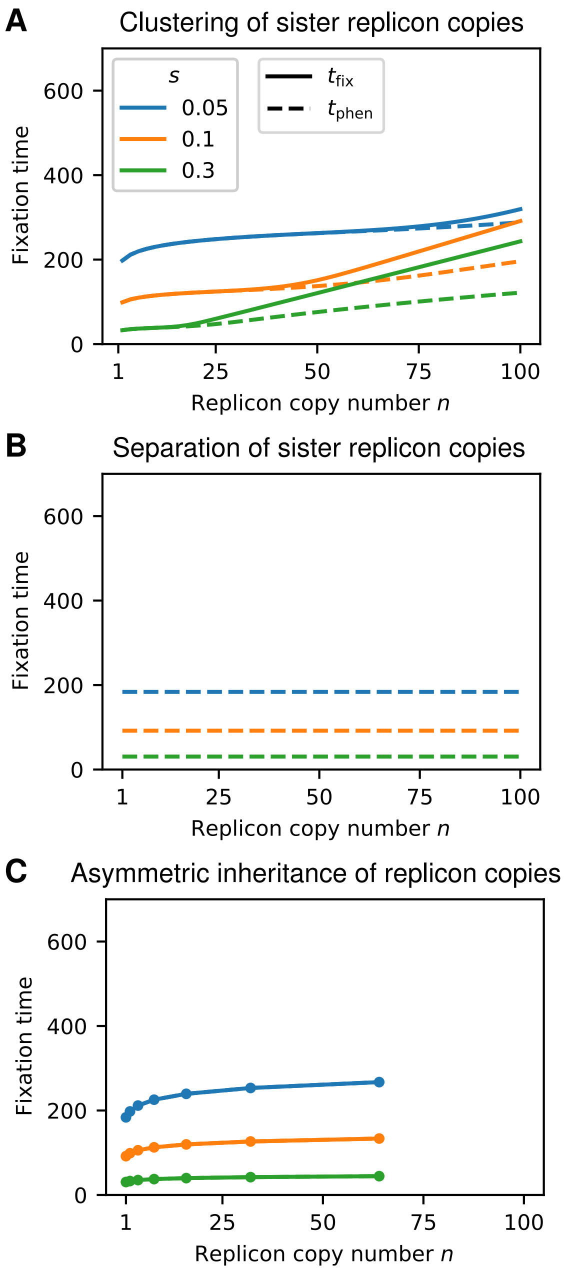


Figure S7: Fixation times as a function of the replicon copy number of simulations for several selection coefficients $s = 0.05$ (blue), 0.1 (orange), 0.3 (green) using the alternative segregation modes. The plots are analogous to Figure 3A (baseline model with *random segregation*) and show results for (A) *clustering of sister replicon copies*, (B) *separation of sister replicon copies*, and (C) *asymmetric inheritance of replicon copies*. Parameter: $f = 0.01$.

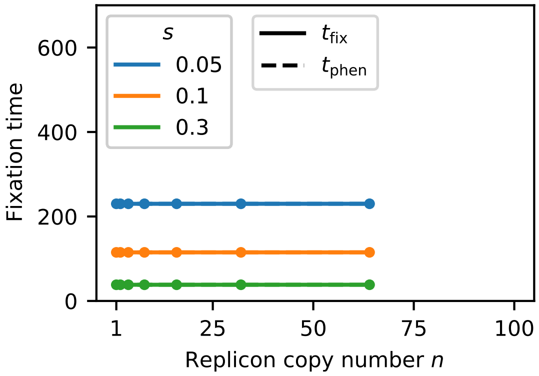


Figure S8: Influence of the replicon copy number $n$ and the strength of selection $s$ on the fixation times for constant initial frequencies of mutant replicon copies $f_{\text{rep}} = f/n = 0.001$ for *asymmetric inheritance of replicon copies*. The figure is analogous to Figure S4A.

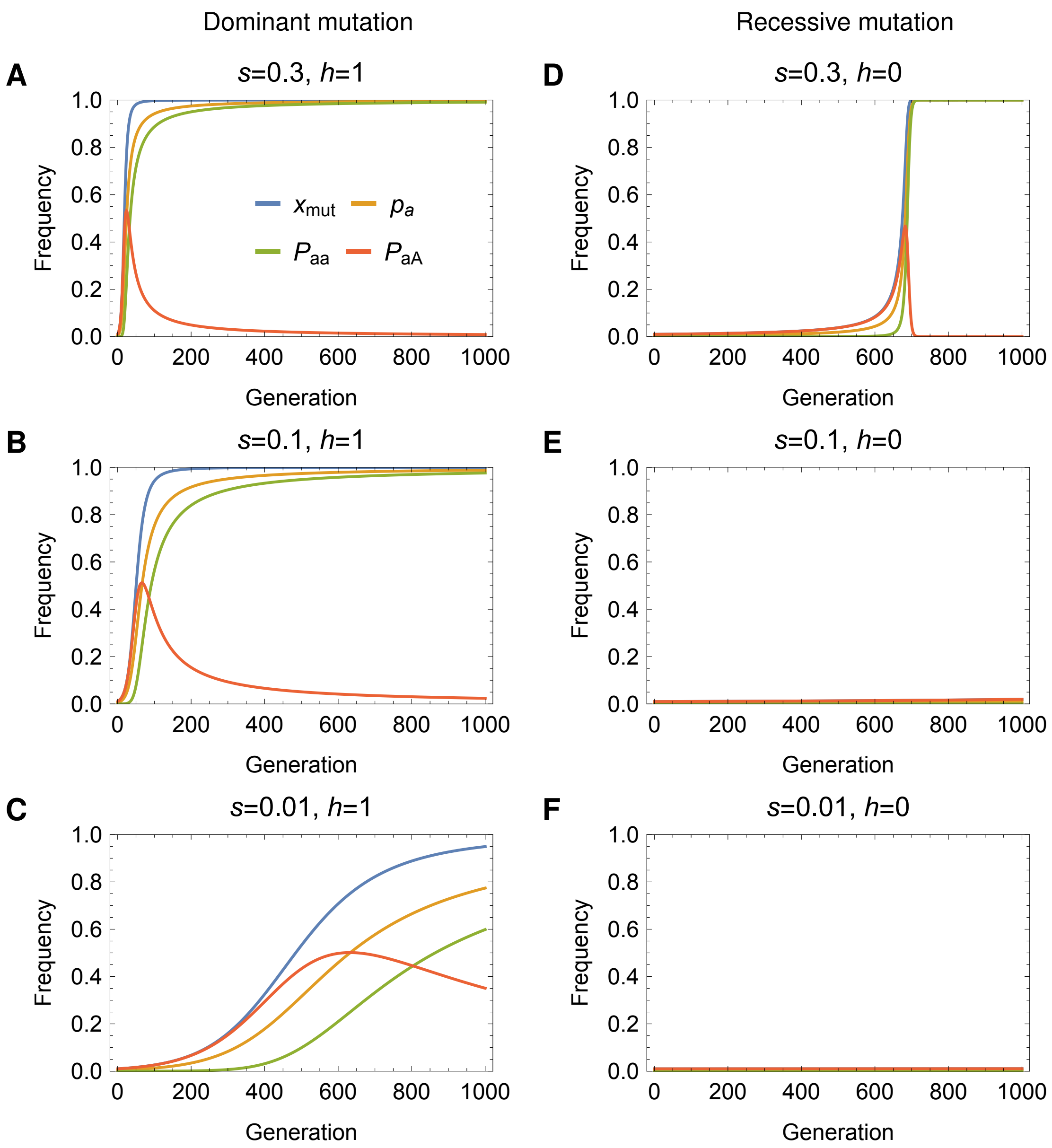


Figure S9: The spread of a beneficial dominant allele (Panels A-D, $h = 1$) and of a beneficial recessive allele (Panels E-H, $h = 0$) in a sexually reproducing population with random mating. The dynamics are modeled by a deterministic Wright-Fisher model with selection (Etheridge, 2011). The variable $p_a$ denotes the frequency of the beneficial allele $a$ in the population. Genotype frequencies of homozygous mutant and heterozygous cells in the subsequent generation are given by $P_{aa} = \frac{p_a^2(1+s)}{\bar{w}}$ and $P_{aA} = \frac{2p_a(1-p_a)(1+hs)}{\bar{w}}$ respectively, where $\bar{w} = 1 + sp_a^2 + 2hsp_a(1 - p_a)$. Phenotypically mutant cells have a fraction $x_{\text{mut}} = P_{aa} + P_{aA}$ for dominant mutations and $x_{\text{mut}} = P_{aa}$ for recessive mutations in the population. The allele frequency is given by $p_a = P_{aa} + P_{aA}/2$. The mutant allele frequency at generation 0 is set to $p_a = f/2$ with $f = 0.01$.

### References

Alison Etheridge. *Some Mathematical Models from Population Genetics*. Springer, 2011.

Wolfram Research, Inc. Mathematica, Version 12.0.0. URL `https://www.wolfram.com/mathematica`. Champaign, IL, 2021.

Eberhard Zeidler. *Oxford Users' Guide to Mathematics*. OUP Oxford, New York, London, 1 edition, 2013.
