## Supplementary material for "Fixation dynamics of beneficial alleles in prokaryotic polyploid chromosomes and plasmids": File S2: balance.py

```
# methods of appendix 4

import numpy as np
import det

# function to calculate the transformation-selection bal.


def balance(n, s, tau, mode="reg", tmax=1e9):

    # function to determine vector of mutational input by transformation
    def tau_omega(x):
        tom = np.zeros(n + 1)
        tom[0] = tau * (-x[0])
        for j in range(1, n + 1):
            tom[j] = tau * ((x[j - 1] - x[j]) * (1 - j / n) + x[j - 1] / n)
        return tom

    sol = det.det(
        n,
        s,
        f=0,
        mode=mode,
        tmax=tmax,
        stopattfix=False,
        stopateq=True,
        tau_omega=tau_omega,
    )

    # raise exception if equilibrium was not found
    if len(sol.t_events[2]) < 1:
        raise Exception(
            "Equilibrium was not found. Check parameters and increase tmax."
        )

    return sol.y.T[-1]
```
