## Supplementary material for "Fixation dynamics of beneficial alleles in prokaryotic polyploid chromosomes and plasmids": File S2: det.py

```
#!/usr/bin/env python3
# -*- coding: utf-8 -*-

"""
    Implementation of methods described in appendix 2 
    (Derivation of the deterministic dynamics) 

    "Fixation dynamics of beneficial alleles on multicopy replicons in prokaryotes"
    bioRxiv (2021) by Santer et al.

    @author: Mario Santer
    @
"""

import numpy as np
from model import p
from scipy.integrate import solve_ivp

import warnings
warnings.filterwarnings("error")

def det(
    n,
    s,
    f=0.01,
    mode="reg",
    tmax=1e9,
    xthr=0.99,
    stopattfix=True,
    stopateq=False,
    tau_omega=lambda x: 0,
):
    """
    simulation of the deterministic dynamics
    """
    # function p^{(1)}_{i->j} and p^{(2)}_{i->j}
    def p1(i, j):
        range1 = [jd for jd in range(n + 1) if jd != j]
        return np.sum([p(i, j, jd, n, mode) + p(i, jd, j, n, mode) for jd in range1])

    def p2(i, j):
        return p(i, j, j, n, mode)

    # matrices (p^{(1)}_{i->j})_ij and (p^{(2)}_{i->j})_ij
    p1M = np.zeros([n + 1, n + 1])
    for i in range(n + 1):
        for j in range(n + 1):
            p1M[i, j] = p1(i, j)
    p2M = np.zeros([n + 1, n + 1])
    for i in range(n + 1):
        for j in range(n + 1):
            p2M[i, j] = p2(i, j)

    # array of replication rates lambda
    l = np.array([1.0 + (i != 0) * s for i in range(n + 1)])
    # linear part of differential equation system
    M1 = np.array(
        [
            [l[i] * (2 * p2M[i, j] + p1M[i, j]) - l[j] * (j == i) for i in range(n + 1)]
            for j in range(n + 1)
        ]
    )

    # non-linear part of differential equation system
    def M2(X):
        return -np.outer(X, l)

    # right-hand side of differential equation (system)
    # tau_omega extension by transformation (see appendix 4)
    def rhs(t, X):
        return (M1 + M2(X)) @ X + tau_omega(X)

    # define the initial state X0 where x0=1-f of cells is wild-type
    pop = np.append([1 - f, f], np.zeros(n - 1))

    # define functions for fixation of phenotype mutants and of homozygous mutant cells
    def phenfix(t, X):
        return X[0] - (1 - xthr)

    def homfix(t, X):
        return xthr - X[n]

    homfix.terminal = stopattfix

    # function to determine if dynamics reached an equilibrium (see appendix 4)
    def hit_eq(t, X):
        nonzero = np.where(X != 0)
        delta = np.max(np.abs(np.divide(rhs(t, X)[nonzero], X[nonzero]))) - 1e-8
        return delta

    hit_eq.terminal = stopateq

    # integrate system
    sol = solve_ivp(
        rhs,
        [0, tmax],
        pop,
        events=[phenfix, homfix, hit_eq],
        method="Radau",
        rtol=1e-5,
        atol=1e-15,
        dense_output=True,
    )

    return sol
```
