## Supplementary material for "Fixation dynamics of beneficial alleles in prokaryotic polyploid chromosomes and plasmids": File S2: model.py

```
#!/usr/bin/env python3
# -*- coding: utf-8 -*-

"""
    Implementation of methods described in appendix 1 (Mathematical formulation 
    of the model and stochastic computer simulations) 

    "Fixation dynamics of beneficial alleles on multicopy replicons in prokaryotes"
    bioRxiv (2021) by Santer et al.

    @author: Mario Santer
    @
"""

import numpy as np
from numpy import eye as e
import math
from scipy.special import binom
from scipy.special import beta
from scipy.stats import betabinom
from warnings import warn
from numpy.random import default_rng
import pickle


def p(i, j1, j2, n, mode="reg"):
    """
    # function p_i->j1j2 for all modes of replication and segregation
    """
    if mode == "reg":  # regular replication and random segregation (baseline model)
        return (
            binom(2 * i, j1)
            * binom(2 * n - 2 * i, n - j1)
            / binom(2 * n, n)
            * (2 * i == j1 + j2)
        )
    elif mode == "ran":  # random replication and random segregation
        p = 0
        for k in range(i, i + n + 1):
            if i>0 and i<n:
                p1 = (
                    binom(n, k - i)
                    * beta(k, 2 * n - k)
                    / beta(i, n - i)
                    * binom(k, j1)
                    * binom(2 * n - k, n - j1)
                    / binom(2 * n, n)
                    * (k == j1 + j2)
                )
            else:
                p1=float(k==2*i)
            p += p1
        return p
    elif mode == "cou":  # clustered segregation of sister replicons
        if n % 2:
            raise Exception("copy number must be even")
        return (
            ((j1 - 1) % 2)
            * binom(i, j1 / 2)
            * binom(n - i, n / 2 - j1 / 2)
            / binom(n, n / 2)
        )
    elif mode == "par":  # separation of sister replicons
        return (i == j1) * (i == j2)
    elif mode == "asy":  # asymmetric inheritance of replicon copies
        if np.modf(np.log(n) / np.log(2))[0]:
            raise Exception("copy number must power of 2")
        if 0 < i and i < n:
            return (2 * i == j1) * (0 == j2)
        elif i == n:
            return (n == j1) * (n == j2)
        elif i == 0:
            return (0 == j1) * (0 == j2)
        else:
            raise Exception("i out of range")
    else:
        raise Exception("mode not valid")


# functions for stochastic simulation


class transitionClass:
    """
    class to control transitions (reactions) and rates in stochastic simulation
    """

    # e_ij1j2k vector (add (delete) individuals of type j1,j2 (i,k))
    def e1(self, i, j1, j2, k):
        return (
            -e(self.n + 1)[i] - e(self.n + 1)[k] + e(self.n + 1)[j1] + e(self.n + 1)[j2]
        )

    def __init__(self, n, s, mode, Nvec):
        self.n = n
        self.ra = range(n + 1)  # range of all types
        self.s = s
        self.l = np.array(
            [1 + s * (i > 0) for i in self.ra]
        )  # array for replication rates
        self.mode = mode
        self.N = sum(Nvec)  # Nveculation size
        # array of factors lambda*p_ij1j2/(N-1) f.a. i,j1,j2
        self.lp = np.zeros((n + 1, n + 1, n + 1))
        for i in range(self.n + 1):
            for j1 in range(self.n + 1):
                for j2 in range(self.n + 1):
                    self.lp[i, j1, j2] += self.l[i] * p(i, j1, j2, n, mode=mode) / (self.N - 1)
        # all reactions
        self.reactionindices = np.array(
            [
                np.array([i, j1, j2, k])
                for i in self.ra
                for j1 in self.ra
                for j2 in self.ra
                for k in self.ra
            ]
        )
        # list of indices of reactions with nonzero effect
        nonzeroreactions = [
            (
                # rate not nan
                (not (math.isnan(self.lp[r[0], r[1], r[2]])))
                *  # and
                # rate is never going to be != 0
                (self.lp[r[0], r[1], r[2]] != 0)
                *  # and
                # reaction has an effect (nonzero)
                (np.any(self.e1(*r) != 0))
            )
            for r in self.reactionindices
        ]
        # update list of indices for online nonzero reactions
        self.reactionindices = self.reactionindices[nonzeroreactions]
        # list of transition vectors e (reactions) of nonzero reactions
        self.reactions = np.array(
            [self.e1(i, j1, j2, k) for (i, j1, j2, k) in self.reactionindices],
            dtype=int,
        )
        # list of rates corresponding to nonzero reactions
        self.rates = np.array(
            [
                Nvec[i] * (Nvec[k] - (i == k)) * self.lp[i, j1, j2]
                for (i, j1, j2, k) in self.reactionindices
            ]
        )

    def update(self, Nvec, mu):
        """
        updates the rates of reactions
        """
        if mu != None:  # mu (last reaction) = None at first reaction, no reaction yet
            # determine reaction that have a changed in rate:
            # check which reactions may have changed in rate since their
            # reaction index i (0) or k (3) are affected by the change in cell-
            # type abundance by the last reaction self.reactions[mu].
            isReactionRateChanged = np.any(
                np.isin(
                    self.reactionindices[:, [0, 3]],
                    np.arange(self.n + 1)[self.reactions[mu] != 0],
                ),
                axis=-1,
            )
            # change rates only of those reactions
            for index in np.arange(len(self.reactionindices))[isReactionRateChanged]:
                (i, j1, j2, k) = self.reactionindices[index]
                self.rates[index] = Nvec[i] * (Nvec[k] - (k == i)) * self.lp[i, j1, j2]
        return self.reactions, self.rates


def stoch(
    n,
    s,
    f=0.01,
    N=1e3,
    mode="reg",
    seed=0,
    tmax=1e9,
    transition=None,
    filename_loadtransition=None,
    filename_savetransition=None,
):
    """
    Runs stochastic simulation with a gillespie algorithm.
    """
    rng = default_rng(seed=seed)

    t = 0
    if np.modf(N * f)[0]:
        warn("cell-type numbers rounded due to the choice of N,f")
    # initial composition of the population
    Nvec = np.array(
        [N - np.round(f * N), np.round(f * N)] + [0 for i in range(2, n + 1)], dtype=int
    )

    t_ts = [t]  # series of time points
    Nvec_ts = [np.copy(Nvec)]  # series of population states

    if filename_loadtransition != None:
        # load transitions from file (for efficient stochastic simulation)
        with open(filename_loadtransition, "rb") as file:
            transition = pickle.load(file)
    else:
        #  new instance of transitionClass to obtain transitions (reactions)
        transition = transitionClass(n, s, mode, Nvec)
        if filename_savetransition != None:  # tmax=0 can be used as a flag to save
            with open(filename_savetransition, "wb") as file:
                pickle.dump(transition, file)

    mu = None

    while True:

        transitions, rates = transition.update(Nvec, mu)
        a0 = sum(rates)
        if a0 == 0:  # absorption
            t_ts.append(t)
            Nvec_ts.append(np.copy(Nvec))
            break

        tnew = t + rng.exponential(scale=1.0 / a0)
        if tnew < tmax:
            mu = rng.choice(len(rates), p=rates / a0)
            t = tnew
            Nvec += transitions[mu]

            t_ts.append(t)
            Nvec_ts.append(np.copy(Nvec))
        else:
            # time-limit reached
            break

    return t_ts, Nvec_ts
```
