## Supplementary material for "Fixation dynamics of beneficial alleles in prokaryotic polyploid chromosomes and plasmids": File S2: Notebook.pdf

### Supplemental material (File S2) for *Fixation dynamics of beneficial alleles in prokaryotic polyploid chromosomes and plasmids* (Santer et al., bioRxiv, 2021)

In[•]:=
```
ClearAll["Global`*"]
$Assumptions = {_ ∈ Reals, n > 1}
```

Out[•]= $\{\_ \in \mathbb{R},\ n > 1\}$

#### Solution of $x_{\mathrm{mut}}(t)$

In[•]:=
```
sol = DSolve[x'[t] == -s * x[t]^2 + x[t] (s + a * (b * Exp[b * t])/(a * Exp[b * t] + (b - a))) && x[0] == f, x, t]
```

Out[•]= $$\left\{\left\{x \to \mathrm{Function}\left[\{t\},\ \frac{e^{s t} \left(-a + b + a e^{b t}\right) f\ (b + s)}{b^2 + a b f - b^2 f - a b e^{s t} f + b^2 e^{s t} f + b s - b f s - a e^{s t} f s + b e^{s t} f s + a e^{(b+s) t} f s}\right]\right\}\right\}$$

(There is only a single solution (as expected).)

In[•]:=
```
xmut[t_] = FullSimplify[x[t] /. sol[[1]]]
```

Out[•]= $$\frac{e^{s t} \left(b + a \left(-1 + e^{b t}\right)\right) f\ (b + s)}{b\ (b + a f - b f + s - f s) + e^{s t} f \left(a e^{b t} s - (a - b)\ (b + s)\right)}$$

Inserting *a, b gives*

In[•]:=
```
ab = {a → (1 + s) (ξ + κ - 1), b → (1 + s) (ξ - 1)};
```

In[•]:=
```
xmut[t_] = FullSimplify[xmut[t] /. ab]
```

Out[•]= $$\left(e^{s t} f\ (-1 + \xi + s \xi) \left(\kappa - e^{(1+s)\, t\, (-1+\xi)}\ (-1 + \kappa + \xi)\right)\right) \Big/ \left(\left(-1 + e^{s t}\right) f \kappa\ (-1 + \xi) - (-1 + \xi)\ (-1 + \xi + s \xi) + f s \left(-1 + \kappa + \xi + \left(-1 + e^{s t}\right) \kappa \xi - e^{t\,(-1+\xi+s\xi)}\ (-1 + \kappa + \xi)\right)\right)$$

#### Decay of wild-type carrying cells $x_0(t)$ and convergence of $\frac{x_{\mathrm{wt}}(t)}{x_0(t)}$

Define $x_{\mathrm{hf}}$. For large t, $\frac{1}{t} \log x_{\mathrm{wt}}(t)$ converges to (1+s) (-1+$\xi$)

In[•]:= $\xi\kappa = \left\{\xi \to \frac{2\,n-3}{2\,n-1},\ \kappa \to \frac{1}{2\,n-1}\right\};$

```
xhf[t_] = FullSimplify[(b * e^(b * t))/(a * e^(b * t) + b - a) /. ab]
xwt[t_] = FullSimplify[1 - xmut[t] * (1 - xhf[t])]
Assuming[{s > 0, (1 + s) ξ > 1, ξ > 0, ξ < 1, f > 0, f < 1, κ < 1 - ξ, κ > 0},
 Limit[1/t Log[xwt[t]], t → ∞]]
```

Out[•]= $\frac{e^{(1+s)\,t\,(-1+\xi)}\,(-1+\xi)}{-\kappa + e^{(1+s)\,t\,(-1+\xi)}\,(-1+\kappa+\xi)}$

Out[•]= $\left((-1+\xi)\left(-1-\left(-1+e^{t\,(-1+\xi+s\,\xi)}\right) f\,(s\,(-1+\kappa)+\kappa)+\xi+s\,\xi\right)\right) \Big/ \left((-1+\xi)\,(-1+\xi+s\,\xi) + f\left(-\left(-1+e^{s\,t}\right)\kappa\,(-1+\xi)+e^{t\,(-1+\xi+s\,\xi)}\,s\,(-1+\kappa+\xi)-s\left(-1+\kappa+\xi+\left(-1+e^{s\,t}\right)\kappa\,\xi\right)\right)\right)$

Out[•]= ConditionalExpression[ $(1+s)\,(-1+\xi)$ , $2\,(1+s)\,\xi > 2+s$ ]

The condition 2 (1+s) $\xi$>2+s must hold to obtain the analytical result (above) with MMA:

For large t, $\frac{x_{wt}(t)}{x_0(t)}$ converges to

```
Assuming[{s > 0, (1 + s) ξ > 1, ξ > 0, ξ < 1, f > 0, f < 1, κ < 1 - ξ, κ > 0},
 Limit[(1 - xmut[t] * (1 - xhf[t]))/(1 - xmut[t]), t → ∞]]
```

Out[•]= $\frac{s\,(-1+\kappa)+\kappa}{(1+s)\,(-1+\kappa+\xi)}$

#### Convergence

The plot below shows an example for *n=32* that Limit[ $\frac{1}{t}$ Log[xwt[t]], t → ∞] equals (1+s)(-1+$\xi$) also if (1+s)$\xi$>1 ⟺ $s > \frac{1}{n-3/2}$ (left vertical line) holds and 2 (1+s) $\xi$>2+s $s > \frac{2}{n-5/2}$ (right vertical line) is not fulfilled.

In[•]:=
```
nrul = {n → 16};
slist = {0.01, 0.02, 0.03, 0.04, 0.05,
   0.06, 0.07, 0.08, 0.09, 0.1, 0.15, 0.2, 0.25, 0.3}
(1 + slist) * ξ /. ξ → (2 n - 3)/(2 n - 1) /. nrul
2 (1 + slist)/(2 + slist) ξ /. ξ → (2 n - 3)/(2 n - 1) /. nrul

y1 = Table[Limit[
    1/t Log[xwt[t]] /. {ξ → (2 n - 3)/(2 n - 1), κ → 1/(2 n - 1), f → 0.01} /. nrul, t → ∞], {s, slist}]
y2 = Table[(1 + s) (-1 + ξ) /. ξ → (2 n - 3)/(2 n - 1) /. nrul, {s, slist}]
ListPlot[{Transpose[{slist, y1}], Transpose[{slist, y2}]},
 PlotMarkers → "OpenMarkers",
 GridLines → {{2/(n - 5/2) /. nrul, 1/(n - 3/2) /. nrul}, {0, 0}},
 AxesLabel → {s, "Limit[1/t Log[xwt[t]],t→∞]"}, PlotRange → Full,
 PlotLegends → {"Numerical solution", (1 + s) (-1 + ξ)}]
```

Out[•]= {0.01, 0.02, 0.03, 0.04, 0.05, 0.06, 0.07, 0.08, 0.09, 0.1, 0.15, 0.2, 0.25, 0.3}

Out[•]= {0.944839, 0.954194, 0.963548, 0.972903, 0.982258, 0.991613, 1.00097, 1.01032, 1.01968, 1.02903, 1.07581, 1.12258, 1.16935, 1.21613}

Out[•]= {0.940138, 0.944746, 0.949309, 0.953827, 0.958301, 0.962731, 0.967119, 0.971464, 0.975768, 0.980031, 1.00075, 1.02053, 1.03943, 1.0575}

Out[•]= {-0.01, -0.02, -0.03, -0.04, -0.05, -0.06, -0.0690323, -0.0696774, -0.0703226, -0.0709677, -0.0741935, -0.0774194, -0.0806452, -0.083871}

Out[•]= {-0.0651613, -0.0658065, -0.0664516, -0.0670968, -0.0677419, -0.0683871, -0.0690323, -0.0696774, -0.0703226, -0.0709677, -0.0741935, -0.0774194, -0.0806452, -0.083871}

Out[•]=
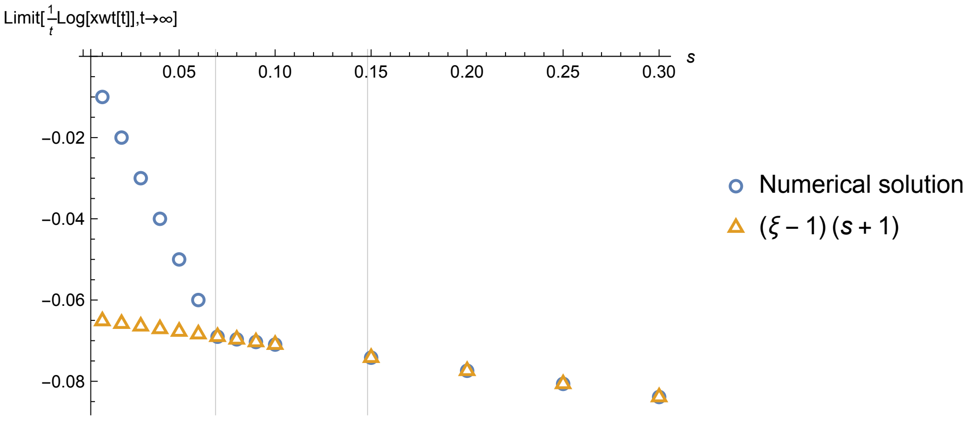

#### Computation of the heterozygosity window

In[∘]:=

```
Hw[n_, s_] = Simplify[-Log[(s (-1+κ)+κ)/((1+s) (-1+κ+ξ))]/((1 + s) (ξ - 1)) /. ξ → (2 n - 3)/(2 n - 1) /. κ → 1/(2 n - 1)]
```

Out[∘]=

$$\frac{(-1 + 2\, n)\, \mathrm{Log}\left[\frac{-1+2\,(-1+n)\, s}{1+s}\right]}{2\,(1 + s)}$$

In[∘]:=

```
Plot[Hw[n, 0.3], {n, 1, 100}]
```

Out[∘]=

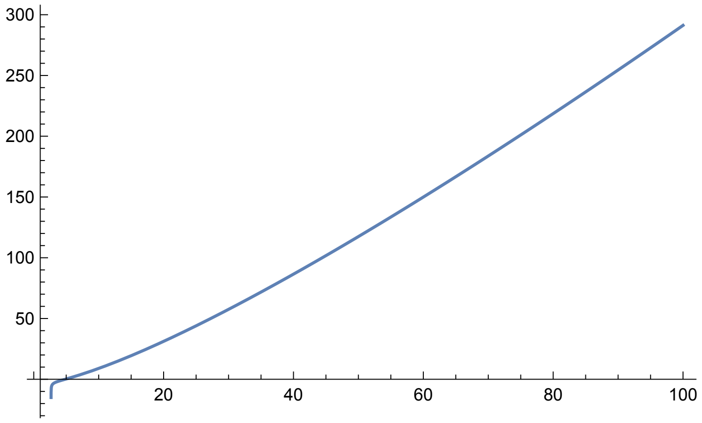
