## Supplementary material for "Fixation dynamics of beneficial alleles in prokaryotic polyploid chromosomes and plasmids": File S2: readme.md

# File S2 – Fixation dynamics of beneficial alleles in prokaryotic polyploid chromosomes and plasmids (bioRxiv 2021)

This supplemental file includes the simulation code repository (mcrfix) and the Mathematica™ notebook (mma) for the manuscript.


Authors: Mario Santer, Anne Kupczok, Tal Dagan, and Hildegard Uecker

## Description of the project

Theoretical population genetics has been mostly developed for sexually reproducing diploid and for monoploid (haploid) organisms, focusing on eukaryotes. The evolution of bacteria and archaea is often studied by models for the allele dynamics in monoploid populations. However, many prokaryotic organisms harbor multicopy replicons --, chromosomes and plasmids -- and theory for the allele dynamics in populations of polyploid prokaryotes remains lacking. Here we present a population genetics model for replicons with multiple copies in the cell. Using this model, we characterize the fixation process of a dominant beneficial mutation at two levels: the phenotype and the genotype. Our results show that, depending on the mode of replication and segregation, the fixation time of mutant phenotypes may precede the genotypic fixation time by many generations; we term this time interval the heterozygosity window. We furthermore derive concise analytical expressions for the occurrence and length of the heterozygosity window, showing that it emerges if the copy number is high and selection strong. Replicon ploidy thus allows for the maintenance of genetic variation following phenotypic adaptation and consequently for reversibility in adaptation to fluctuating environmental conditions.

## mcrfix/

The repository contains python source code (src).

The source code is structured according to the methods described in the appendix of the manuscript.

##### Modules

```model.py``` contains two functions:

-  ```p(i,j1,j2,n,mode)``` to calculate $p_{i\rightarrow j_1 j_2}$ for all modes of replication and segregation (parameter ```mode```) described in the main text. This function gets used by both function for stochastic and deterministic simulations.
- ```stoch``` for stochastic simulations. The function also makes use of the defined class ```transitions``` that computes the transition vectors and corresponding rates between population states. The main parameters are ```n``` (replicon copy number), ```s``` (strength of selection), ```f``` (initial frequency of mutant cells), ```N``` (population size), and ```mode``` (as above). An instance of ```transitions``` can be loaded from a file and should be used for efficiency reasons. 

```det.py``` contains a function to run deterministic simulations. Main parameters are the almost the same as in ```stoch``` (above, population size is obviously not a parameter here). Moreover, there is a parameter  ```xthr```for the threshold of fixation (see main text).

- To solve the determinstic dynamics we use a matrix equation reflecting the system of ordinary differential equations (see main text):
  $$
  \begin{aligned}
  \frac{\mathrm{d}x_j}{\mathrm{d}t} &=
  \sum_{i=0}^{n} \left\{ x_i \lambda_i (2 p^{(2)}_{i \rightarrow j} +  p^{(1)}_{i \rightarrow j} - x_j) \right\}
  - x_j \lambda_j \\
  &= \sum_{i=0}^{n} \lambda_i (2 p^{(2)}_{i \rightarrow j} +  p^{(1)}_{i \rightarrow j}) x_i - \lambda_i \delta_{ij} x_i + \sum_{i=0}^{n} - \lambda_i x_j x_i\\
  &= \sum_{i=0}^{n} \lambda_i (2 p^{(2)}_{i \rightarrow j} +  p^{(1)}_{i \rightarrow j} - \delta_{ij}) x_i - \sum_{i=0}^{n} (\mathbf{x} \otimes \mathbf{\lambda})_{ji} x_i\\
  \Leftrightarrow \frac{\mathrm{d}\mathbf{x}}{\mathrm{d}t}
  &= \underbrace{(\lambda_i (2 p^{(2)}_{i \rightarrow j} +  p^{(1)}_{i \rightarrow j} - \delta_{ij}))_{ji\in {0,\dots,n}}}_{\mathtt{M1}} \mathbf{x} + \underbrace{(\mathbf{x} \otimes \mathbf{\lambda})}_{ \mathtt{M2}(\mathbf{x})} \mathbf{x}\\
  & = (\mathtt{M1}+\mathtt{M2}(\mathbf{x}))\mathbf{x}
  \eqqcolon \mathtt{rhs(t,\mathbf{x})}
  \end{aligned}
  $$
  

  $\mathbf{x}=(x_0,\dots,x_n)$, $\mathbf{\lambda}=(\lambda_0,\dots,\lambda_n)$, $\delta$ denotes Kronecker's delta, $\otimes$ is the tensor (outer) product.

```balance.py``` contains a function ```balance``` to obtain the transformation-selection balance. The main parameters are ```n```, ```s```, ```tau``` where the latter is the transformation rate.

### Installation

Please make sure you have installed the following python packages :

1. [numpy](https://numpy.org/)
2. [scipy](https://www.scipy.org/)

### Contributing

The source code of ```mcrfix``` is also stored on ```github.com/mariosanter/mcrfix```. Please consider a contribution or contact Mario Santer, email: santer{at}evolbio.mpg.de.

## mma/

This folder contains a notebook including computer algebra methods referred to in File S2. 

## License

```mcrfix/``` [MIT](https://choosealicense.com/licenses/mit/)

```mma/``` [MIT](https://choosealicense.com/licenses/mit/) 
